# scaDA: A Novel Statistical Method for Differential Analysis of Single-Cell Chromatin Accessibility Sequencing Data

**DOI:** 10.1101/2024.01.21.576570

**Authors:** Fengdi Zhao, Xin Ma, Bing Yao, Li Chen

## Abstract

Single-cell ATAC-seq sequencing data (scATAC-seq) has been widely used to investigate chromatin accessibility on the single-cell level. One important application of scATAC-seq data analysis is differential chromatin accessibility analysis. However, the data characteristics of scATAC-seq such as excessive zeros and large variability of chromatin accessibility across cells impose a unique challenge for DA analysis. Existing statistical methods focus on detecting the mean difference of the chromatin accessible regions while overlooking the distribution difference. Motivated by real data exploration that distribution difference exists among cell types, we introduce a novel composite statistical test named “scaDA”, which is based on zero-inflated negative binomial model (ZINB), for performing differential distribution analysis of chromatin accessibility by jointly testing the abundance, prevalence and dispersion simultaneously. Benefiting from both dispersion shrinkage and iterative refinement of mean and prevalence parameter estimates, scaDA demonstrates its superiority to both ZINB-based likelihood ratio tests and published methods by achieving the highest power and best FDR control in a comprehensive simulation study. In addition to demonstrating the highest power in three real sc-multiome data analyses, scaDA successfully identifies differentially accessible regions in microglia from sc-multiome data for an Alzheimer ‘s disease (AD) study, regions which are most enriched in GO terms related to neurogenesis, the clinical phenotype of AD, and SNPs identified in AD-associated GWAS.

**Author summary:** Understanding the cis-regulatory elements that control the fundamental gene regulatory process is important to basic biology. scATAC-seq data offers an unprecedented opportunity to investigate chromatin accessibility on the single-cell level and explore cell heterogeneity to reveal the dynamic changes of cis-regulatory elements among different cell types. To understand the dynamic change of gene regulation using scATAC-seq data, differential chromatin (DA) analysis, which is one of the most fundamental analyses for scATAC-seq data, can enable the identification of differentially accessible regions between cell types or between multiple conditions. Subsequently, DA analysis has many applications such as identifying cell type-specific chromatin accessible regions to reveal the cell type-specific gene regulatory program, assessing disease-associated changes in chromatin accessibility to detect potential biomarkers, and linking differentially accessible regions to differentially expressed genes for building a comprehensive gene regulatory map. This paper proposes a novel statistical method named “scaDA” to improve the detection of differentially accessible regions by performing differential distribution analysis. scaDA is believed to benefit the research community of single-cell genomics.

## Introduction

Open chromatin offers permissible physical interactions to transcription factors and cis-regulatory DNA elements (CREs) for cooperatively regulating gene expression in a complex interplay between them [1, 2]. With the advent of next-generation sequencing (NGS) techniques, ATAC-seq has become one of the most popular NGS technologies to profile genome-wide open chromatin regions. Compared to its predecessors such as DNase-seq, ATAC-seq requires much fewer cells and is less laborious in the experimental preparation [2]. However, bulk ATAC-seq cannot determine chromatin accessibility on the single-cell level, which limits its application for identifying cell subpopulations. To overcome these limitations, single-cell ATAC-seq (scATAC-seq) has been developed to measure chromatin accessibility on the single-cell level benefiting from the advantage of ATAC-seq protocol such as obtaining sufficient biological signals while requiring fewer cells. With the advent of scATAC-seq technology, the data richness of single-cell genomics has been significantly expanded. It is straightforward to profile separate scATAC-seq and scRNA-seq data for the same sample to learn the gene regulatory relationship [3]. Nevertheless, the “unmatched” data, where RNA and ATAC are not jointly profiled in the same cells, impose challenges such as batch effect in the data analysis. Recent 10x sc-multiome protocol enables the simultaneous profiling of the two modalities in the same cell, which provides an opportunity to integrate two data modalities for a joint unbiased definition of cellular state.

One important analysis for scATAC-seq data is differential chromatin accessibility (DA) analysis, which aims to identify DA regions between cell types or conditions. The standard bioinformatics pipeline for identifying DA regions usually consists of three steps [4, 5]. First, read mapping and peak calling to identify open chromatin regions, known as “peaks”. Read counts for each peak and each cell are calculated to build a peak-by-cell matrix. Second, cell clustering is performed to identify cell types after a series of data quality control, data normalization, and dimension reduction steps. Lastly, DA analysis can be performed for each peak between two cell types. For scATAC-seq samples from multiple conditions, a unified set of peaks across all samples will be created. After integration and alignment of the samples, DA analysis can be performed for each cell type across multiple conditions. The major difference of different bioinformatics pipelines lies in the steps to create the peaks. Instead of doing one round of peak-calling using all reads, cluster-specific peak calling can be performed [6]. Particularly, DA analysis has three important applications. First, DA regions can be linked to differentially expressed genes for building a comprehensive gene regulatory map. As an example, DA regions have been found localized to promoters, the majority of which are closely associated with differentially expressed genes in the adult human kidney [7]. Second, cell type-specific DA regions enable the identification of different cell subpopulations. For example, DA analysis can distinguish cell types by cluster-specific DA regions associated with cell type-specific marker genes identified from scRNA-seq [8]. Third, DA analysis can detect disease-associated DA regions by comparing disease and control scATAC-seq samples. For example, DA analysis successfully identifies disease-associated DA regions from scATAC-seq data collected from both peripheral venous blood in sporadic amyotrophic lateral sclerosis patients and health controls [9].

Nevertheless, the distinctive characteristics of scATAC-seq data introduce specific challenges in DA analysis. First, the scATAC-seq peak-by-cell count matrix exhibits higher sparsity (about 3% non-zero entries) compared to the scRNA-seq gene-by-cell count matrix (more than 10% non-zeros) due to smaller dimension of genes relative to peaks and lower gene dropout rates [10, 11]. Second, the biological variation, which represents the heterogeneity of chromatin accessibility among cells in the same cell types, is non-negligible, leading to the “overdispersion”. Third, the sample size, i.e. number of cells in rare cell types is small, which may reduce the power of DA analysis. Various statistical methods have been developed for DA analysis. Nonparametric methods, like the Wilcoxon rank-sum test, have been adopted by state-of-the-art scATAC-seq analysis pipelines such as Signac [5] and scATAC-pro [12] due to their robustness to distribution assumptions. However, the Wilcoxon test cannot adjust for covariates, provide interpretable effect sizes, and is generally less powerful for rare cell types. In addition, Signac adopts a logistic regression model as another option for DA analysis, treating condition membership as the covariate and the total number of counts in each cell as a latent variable. Nevertheless, the high sparsity and overdispersion make the logistic regression model less desirable. Moreover, methods developed for differential expression (DE) analysis can be applied for DA analysis directly such as edgeR [13], which is based on the negative binomial test for DE analysis of bulk RNA-seq, and MAST [14] for DE analysis of scRNAseq, which models log2 transformed TPM expression matrix in a two-part generalized regression model. These methods, however, may not be optimal for scATAC-seq data due to the inherent differences in data characteristics between scATAC-seq and scRNA-seq data.

Given the excessive zeros of scATAC-seq data (Fig 2), we perform a real data exploration using a zero-inflated negative binomial (ZINB) model and find the”distribution difference”, where all three parameters are different among cell types. Motivated by this observation, we hereby develop an omnibus test, named “scaDA” (Single-Cell ATAC-seq Differential Chromatin Analysis) based on ZINB model for scATAC-seq DA analysis. The contribution of scaDA mainly lies in two aspects. First, to our best knowledge, scaDA is the first statistical method, which focuses on testing distribution difference in a composite hypothesis, while most existing methods only focus on testing mean difference. The composite hypothesis may be more powerful in the presence of distribution differences. Second, scaDA improves the parameter estimation by leveraging an empirical Bayes approach for dispersion shrinkage and iterative estimation of mean and prevalence parameters. Using a comprehensive simulation design, we demonstrate that scaDA is superior to both ZINB-based likelihood ratio tests and published methods by achieving the highest power and best FDR control. In addition, scaDA is more powerful than published methods in DA analysis of three real sc-multiome data. Furthermore, scaDA successfully identifies AD-associated DA regions by both GO term analysis and GWAS enrichment analysis.

**Figure 1:**
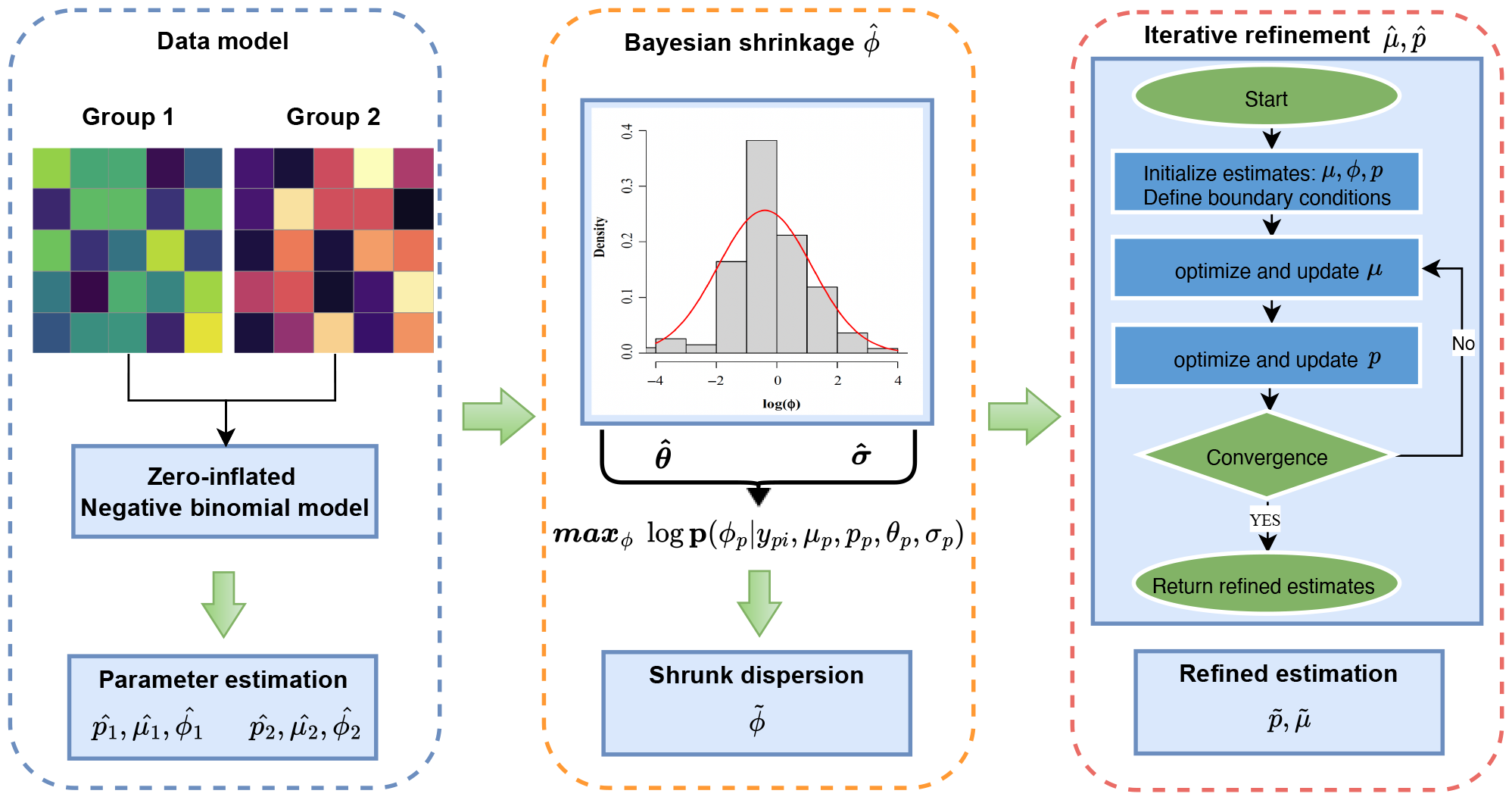
scaDA method overview. scaDA contains three components: Data model, Bayesian shrinkage of 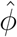, and iterative optimization of 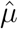 and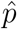

**Figure 2:**
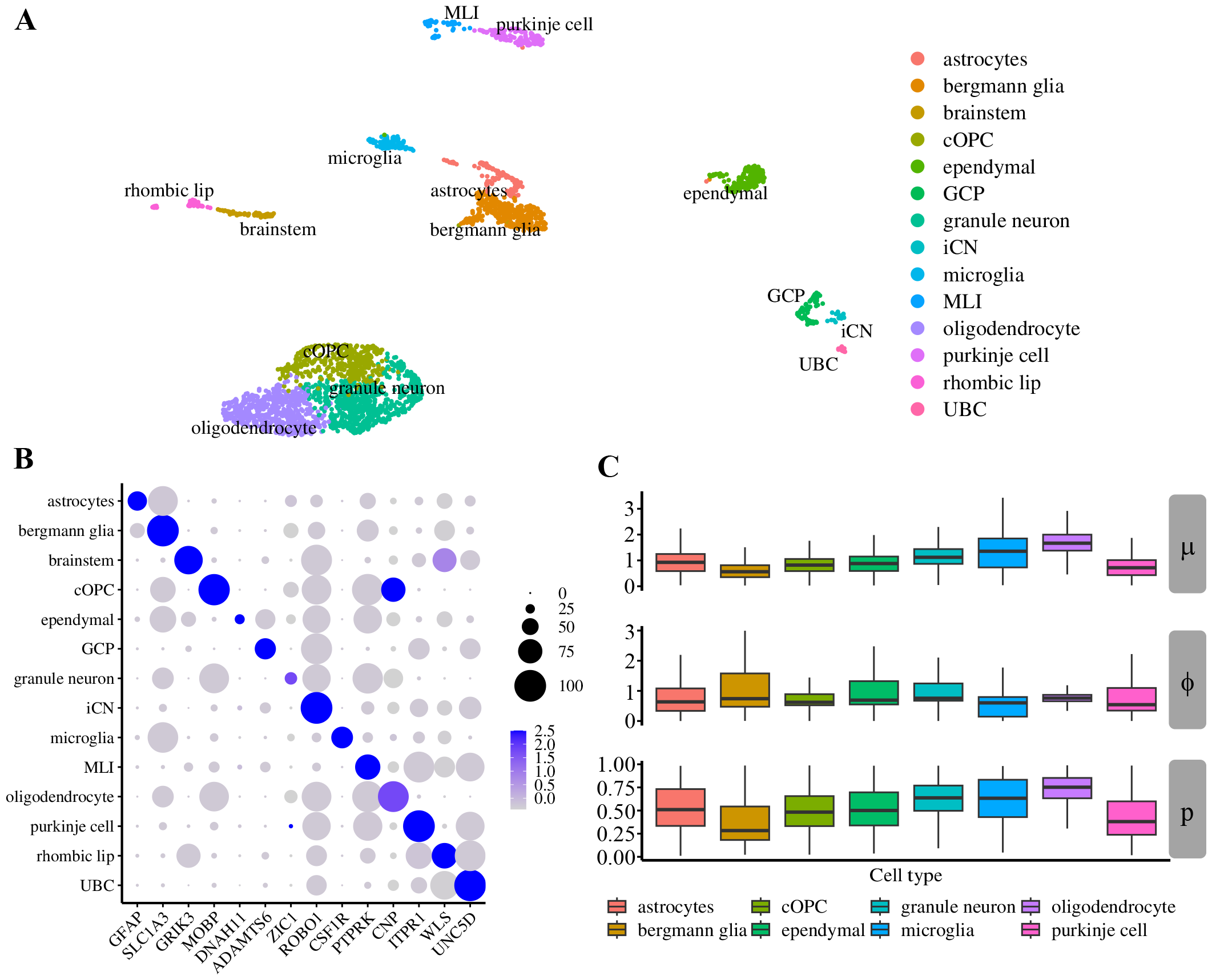
Human Brain 3K analysis. **A**. Clustering analysis of “Human Brain 3K” shows 14 annotated cell types as shown in the UMAP. **B**. Expression of gene markers used for cell type annotation. **C**. Distribution of EM initial estimates for three parameters in ZINB model, which include mean (*μ*), prevalence (*p*), and dispersion (*ϕ*) across all peaks for 8 cell types with more than 100 cells.

## Materials and methods

### scaDA availability

The scaDA has been implemented as an R package and is freely available from github (https://github.com/fzhaouf/scaDA)

### Statistical method

#### Data Model

Without group indicator, we denote *y*_*pi*_ as read count in peak *p* in sample *i* and model *y*_*pi*_ based on Zero-Inflated Negative Binomial (ZINB) model as

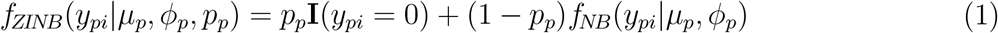

which is a mixture distribution of a zero-inflated component and a negative binomial distribution (NB) by assigning *p*_*p*_ to extra zeros and 1*−p*_*p*_ to the NB distribution *f*_*NB*_(*y*_*pi*_|*μ*_*p*_, *ϕ*_*p*_). *p*_*p*_ (0 *≤ p*_*p*_ *≤* 1) is the prevalence parameter, representing the probability of excess zeros.

Under the Gamma-Poisson parameterization, the NB distribution *f*_*NB*_(*y*_*pi*_|*μ*_*p*_, *ϕ*_*p*_) is formularized as

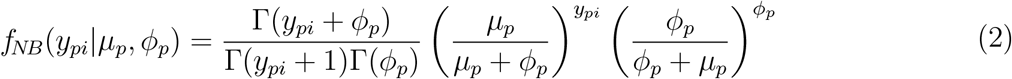

where *μ*_*p*_ and *ϕ*_*p*_ characterize the mean and dispersion of the NB distribution.

Thus, the distribution of read counts in each peak is characterized by its own three parameters *p*_*p*_, *μ*_*p*_ and *ϕ*_*p*_. By inducing the zero-inflated component, scaDA benefits from modeling the excessive zeros, which is more evident in scATAC-seq data. The excessive zeros can either be biological excessive zeros because ATAC signals are truly not expressed in peaks and technical excessive zeros due to dropout events in the sequencing process.

#### Initial estimation of *p, μ* and *ϕ*

Given the observed count data **y**_*p*_ for peak *p*, the likelihood is given by

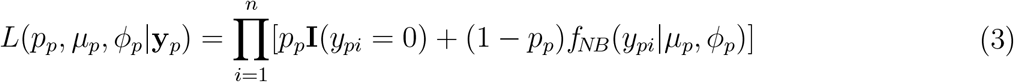

Accordingly, the log-likelihood function is given by

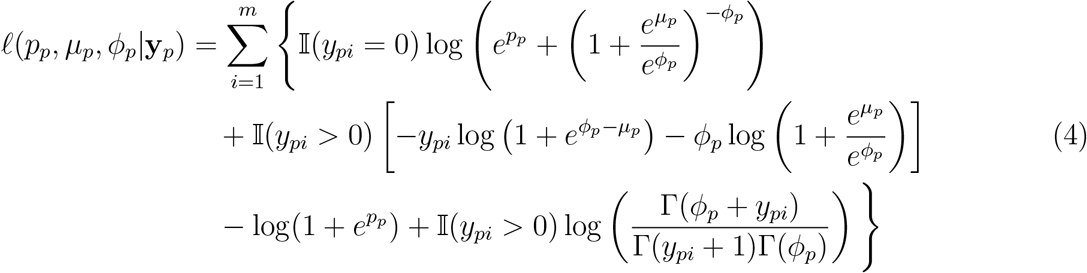

By introducing a latent variable *r*_*pi*_, which indicates whether *y*_*pi*_ is from zero point mass (*r*_*pi*_ = 1) or the NB distribution (*r*_*pi*_ = 0), the maximization of the log-likelihood function is implemented by EM algorithm for a complete likelihood function (Equation 5). In scaDA ‘s implementation, we estimate parameters using the “zeroinfl” function from the R package pscl (Zeileis, 2008).

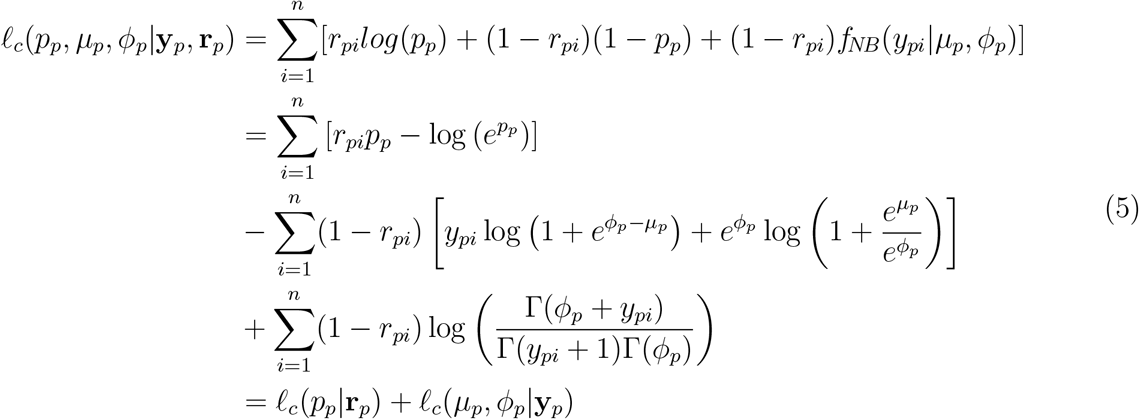

#### Dispersion shrinkage

Accurate estimation of dispersion parameter *ϕ* is crucial for DA analysis to stabilize the variance estimate, especially considering either small numbers of cells or highly sparse read counts. Though multiple previous works improve the estimation of the dispersion parameter for each peak/gene in an empirical Bayes framework by leveraging the estimates of dispersion parameter from all peaks/genes [15, 13, 16], most of them focus on bulk RNA-seq data, which is usually modeled by NB distribution. It is unclear if borrowing information across other peaks will help improve the estimation of *ϕ* for each peak and ultimately improve the power for DA analysis.

We further perform a real data analysis to explore the empirical distribution of initial estimates of *ϕ* for all peaks in Granule neuron of “Human Brain 3K” data. For each peak, we apply the above EM algorithm to estimate the dispersion parameters. As shown in Supplementary **Figure S1**, the genome-wide distribution of logarithm dispersion parameter estimates is approximately Gaussian with mean = 0.03 and SD = 0.57, suggesting that the dispersion parameters can be well-approximated by a log-normal distribution (Supplementary **Figure S1**). This observation motivates us to adopt log-normal distribution as a prior for obtaining a posterior estimates of *ϕ*. Accordingly, the conditional posterior distribution of *ϕ*_*p*_ given observed read counts and other parameters is shown in Eq 6.

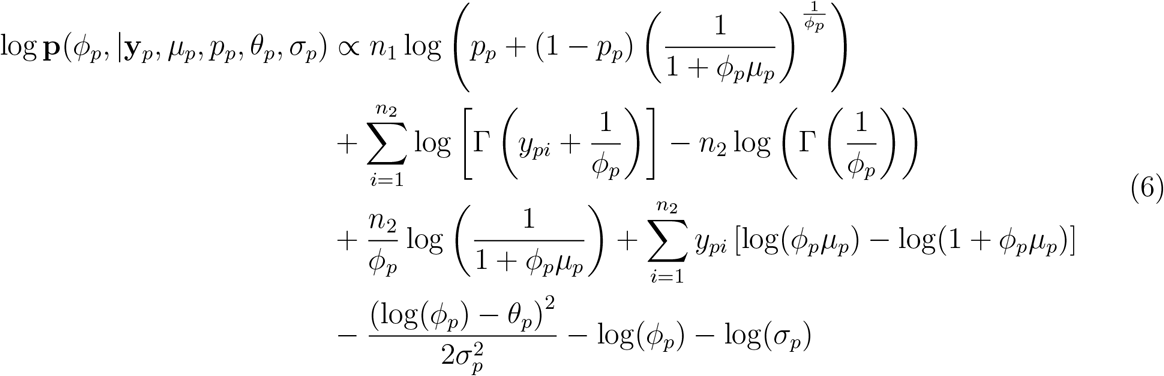

Motivated by Wu et al. [16], instead of using a computationally intensive MCMC approach, we compute the posterior mode by maximizing Eq 6 to obtain a point estimate of *ϕ*_*p*_, denoted As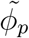.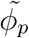is essentially the shrunken estimate of *ϕ*_*p*_ in the empirical Bayes framework. In the conditional posterior distribution, two parameters *p*_*p*_ and *μ*_*p*_ and two hyper-parameters θ and σ need to be determined first before maximizing log **p**(*ϕ*_*p*_, |*y*_*pi*_, *μ*_*p*_, *p*_*p*_, *θ*_*p*_, *σ*_*p*_) to obtain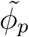. It is straightforward that *p*_*p*_ and *μ*_*p*_ can be obtained from the estimates in the EM algorithm. Two hyper-parameters,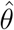 and 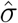in the log-normal distribution can be inferred from dispersion estimates 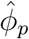 in the EM algorithm across all peaks. Particularly,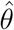 is estimated by the median of log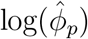 .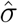 is estimated by using sample variance of 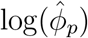 after adjusting the variation due to estimating log(*ϕ*_*p*_) [16]. Afterwards, by plugging in 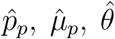 and 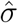, we maximize Eq 6 with respect to *ϕ*_*p*_ using the Newton-Raphson method.

### Iterative optimizing estimates of *μ* and *p*

As the initial estimate 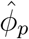 in each peak *j* is optimized into the empirical Bayes shrunken estimate 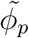 in the last step, the full likelihood for each peak may need to be re-maximized, which needs to re-estimate 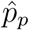 and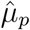 for achieving this purpose. The updated estimates 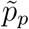 and 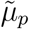, together with 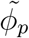 will be treated as the final estimates for each peak. Specifically, we have developed an iterative optimization algorithm in the context of the updated dispersion 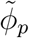, as detailed in Algorithm 1

#### Algorithm 1

Iterative optimizing the estimates of *μ* and *p*

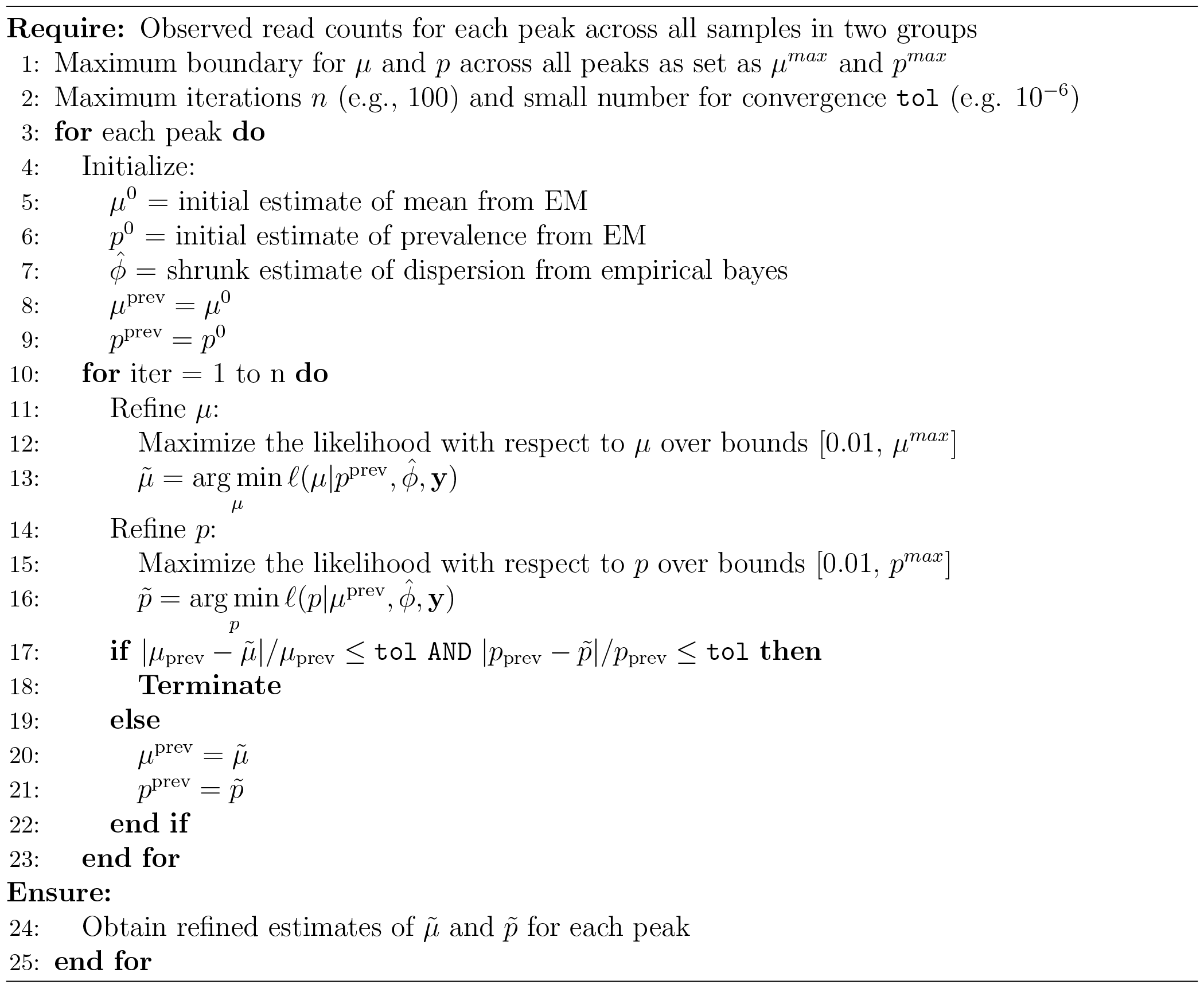

### Composite hypothesis testing

For each peak, we adopt the likelihood ratio test (LRT) to perform the composite hypothesis testing. We denote the number of cells for the two groups as *n*_1_ and *n*_2_ and the corresponding parameters as Θ_1_=(*p*_1_, *μ*_1_, *ϕ*_1_) and Θ_2_=(*p*_2_, *μ*_2_, *ϕ*_2_).

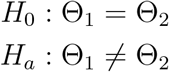

The likelihood ratio statistic is formulated as,

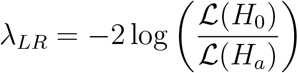

For the reduced model under *H*_0_, all cells from the two groups are pooled together to estimate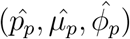 as,

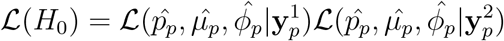

For the full model under *H*_*a*_, 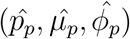 are estimated for each group independently as,

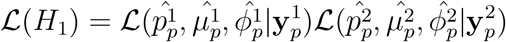

The likelihood ratio statistic *λ*_*LR*_ follows a 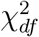 distribution with degree freedom of 3 in this special case and the *p*-value is obtained as

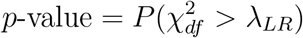

### Evaluated methods

We evaluate scaDA against two sets of methods. The first s et o f c ompared m ethods consists of ZINB(*μ*) testing the mean difference w ith o ne d egree o f f reedom (1 d .f.), Z INB(*p*) testing the prevalence difference w ith o ne d egree o f f reedom (1 d .f.), ZINB(*μ, p*) t esting o n b oth the mean and prevalence difference with two degrees of freedom (2 d .f.), ZINB(*μ, ϕ, p*) testing on the difference o f t hree p arameters s imultaneously w ith t hree d egrees o f f reedom (3 d .f.), which has been adopted for differential d istribution a nalysis i n t he m icrobiome s tudy [17]. T he fi rst se t of methods is compared to evaluate scaDA ‘s advantage in jointly testing three parameters (*μ, p, ϕ*) simultaneously, incorporating the shrunk *ϕ* and iterative refinement of *μ* and *p* estimates.

The second set of methods for comparison includes published approaches like the negative binomial model (NegBin), edgeR [13], scATAC-pro[12], Signac[5] and MAST[14]. Specifically, NegBin is frequently used to model sequencing count data. edgeR, extendding NegBin, adopts an empirical Bayesian method for dispersion shrinkage and is widely used for differential expression (DE) analysis in bulk RNA-seq. MAST, a popular DE analysis method for scRNA-seq data, models the log2 transformed TPM (transcripts per million) expression matrix using a two-part generalized regression framework instead of directly modeling the count data. scATAC-pro and Signac are two methods/pipelines specifically d esigned f or d ifferential ac cessibility (DA) analysis for scATAC-seq data. scATAC-pro adopts the Wilcoxon rank sum test as the default method to perform differential accessibility analysis. Signac uses a logistic regression model as default for DA analysis with condition membership as the covariate and the total number of counts per cell as a latent variable.

### Evaluation metric

After conducting DA analysis, each method calculates a raw p-value for qualifying the distribution difference in each p eak. To control for false positives in multiple testing, the False Discovery Rate (FDR) is calculated using the Benjamini-Hochberg (BH) approach [18]. Performance will be evaluated based on both power (True Positive Rate, TPR) and FDR control. The observed FDR is defined as the expectation of false discovery proportion (FDP). The formulas for TPR and FDP are:

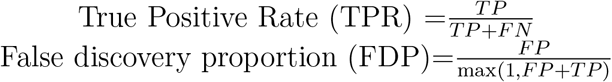

Specifically, TPR is calculated given a nominal FDR level (i.e., 0.05) to claim differential peaks and non-differential peaks. Subsequently, FDR control is measured by the comparison between the nominal FDR level and observed FDP.

### Data description

#### Human Brain 3K

The single-cell multiome data, referred to as “Human Brain 3K”, is generated from the Single Cell Multiome ATAC + Gene Expression Sequencing protocol from 10X Genomics, which profiles gene expression and chromatin accessibility for each cell from frozen healthy human brain cerebellum region. The processed data by Cell Ranger is available on 10X Genomics Datasets, which include 3,233 cells, and 13,629 genes linked to 88,843 peaks.

#### Human PBMC 10K

The single-cell multiome data, referred to as “Human PBMC 10K”, is generated from the Single Cell Multiome ATAC + Gene Expression Sequencing protocol from 10X Genomics, which profiles gene expression and chromatin accessibility for each cell from human peripheral blood mononuclear cells (PBMCs). The processed data by Cell Ranger is available on 10X Genomics Datasets, which include 12,012 cells, and 13,958 genes linked to 79187 peaks.

#### Human AD

The single-cell multiome data, referred to as “Human AD”, consists of a total of 191,890 single-cell nuclei from 20 postmortem samples of Human prefrontal cortex from both AD and cognitively healthy controls [19]. Among the 20 samples, 12 samples are AD, and 8 are healthy controls. Instead of co-profiling gene expression and ATAC in each cell, the RNA and ATAC modalities are profiled in separate scRNA-seq and scATAC-seq datasets. The data is available from the Gene Expression Omnibus (GEO) under the accession number GSE174367.

## Results

### Differential distribution among cell types in “Human Brain 3K”

An important feature of scaDA is the composite hypothesis, which focus on testing the differential distribution among cell types. Without loss of generality, we analyse “Human Brain 3K” to explore the differential distribution among cell types in scATAC data. Specifically, we perform clustering and cell type annotation using gene markers from a published study focused on human cerebellar development [20]. Consequently, 14 cell types have been annotated, which include astrocytes, bergmann glia, brainstream, cOPC, ependymal, GCP, granule neuron, iCN, microglia, MLI, oligodendrocyte, purkinje cell, rhombic lip, UBC (Fig 2A). Gene marker expression across the 14 cell types demonstrates the gene markers can distinguish the cell types clearly (Fig 2B). We further remove GCP, iCN, MLI and UBC with cell numbers less than 100 and retain 8 cell types with more cell numbers for a robust parameter estimation.

Using EM algorithm for initial estimation for ZINB model, we obtain the cell-type specific parameters, which include mean (*μ*), prevalence (*p*), and dispersion (*ϕ*) for all peaks. We observe a large variation of mean, prevalence, and dispersion among cell types, indicating DA analysis should be performed in a cell type-specific manner. Additionally, the distribution difference of the three parameters between two cell types varies in different patterns (Fig 2C). Notably, we notice that the dispersion is lower in cOPC than in granule neurons and the other two parameters are similar, which confirms the necessity to include dispersion as one parameter in the composite hypothesis test. Moreover, differences in the distributions of all three parameters are observed between multiple cell type paires such as astrocytes and bergmann glia, or oligodendrocyte and purkinje cell. These findings highlight the importance of developing a robust composite hypothesis test, which can jointly test the three parameters simultaneously. However, most existing methods focus on testing only the mean difference but ignore the distribution difference of other parameters such as prevalence and dispersion, which is further confirmed by real data exploration. To fill the gap, we hereby develop an omnibus test, named scaDA, to test the distribution difference of all three parameters in a composite hypothesis test (more details in the Method Section). The test will be more powerful when the underlying differential distribution pattern of two cell types is unknown. Accordingly, to demonstrate the robustness of scaDA, we design a comprehensive simulation strategy, where each one of the three parameters will change or three parameters change simultaneously, to highlight the importance of considering the distribution difference.

### Simulation studies

### Overview of simulation design

We evaluate the performance of scaDA along with other methods in a real data-driven simulation design. Without loss of generality, we choose the three parameters estimated in the cell type “Granule neuron” from “Human Brain 3K” (Fig 2C) as the baseline parameters. Specifically, we simulate count data semi-parametrically. In all simulations, the mean and prevalence are randomly sampled in pairs from the estimates of mean and prevalence. The dispersion parameters are generated from the log-normal parametric distribution. The simulation allows for a more realistic assessment of power analysis and FDR control, and fair comparison with existing methods.

For non-differential peaks, we allow the two groups to have the same parameter values as the baseline. For differential peaks, we fix the parameter values in Group 1 and vary the parameter values in Group 2 with different effect sizes of selected parameters to create power curves. In the first three scenarios, we modify one parameter such as mean, prevalence, or dispersion for differential peaks in Group 2 at a time, holding other parameters the same as the baseline. In the fourth scenario, we change all three parameters simultaneously for differential peaks in Group 2. To create parameter values in Group 2, we multiply and divide an effect size in terms of log2 fold change from the baseline values for differential peaks in equal proportion. The log2 fold change of selected parameters is changed from 0.5 to 3.0 with a step size of 0.5. Using the parameter setting in the four scenarios, we assume 20% differential peaks and simulate the read counts based on ZINB for 4000 peaks across 100 cells in each group (Eq 3).

### Power analysis for scaDA and ZINB-based likelihood ratio tests

We perform a simulation study to evaluate the power of scaDA and other ZINB-based likelihood ratio tests under different scenarios when the underlying differential accessibility is driven by one or several parameters across different effect sizes in order to have a better understanding of the model performance. Fig 3 summarizes the power of five methods in each scenario. As expected, the overall power of all methods increases when the effect size of parameter difference (i.e. log2 fold change) increases.

**Figure 3:**
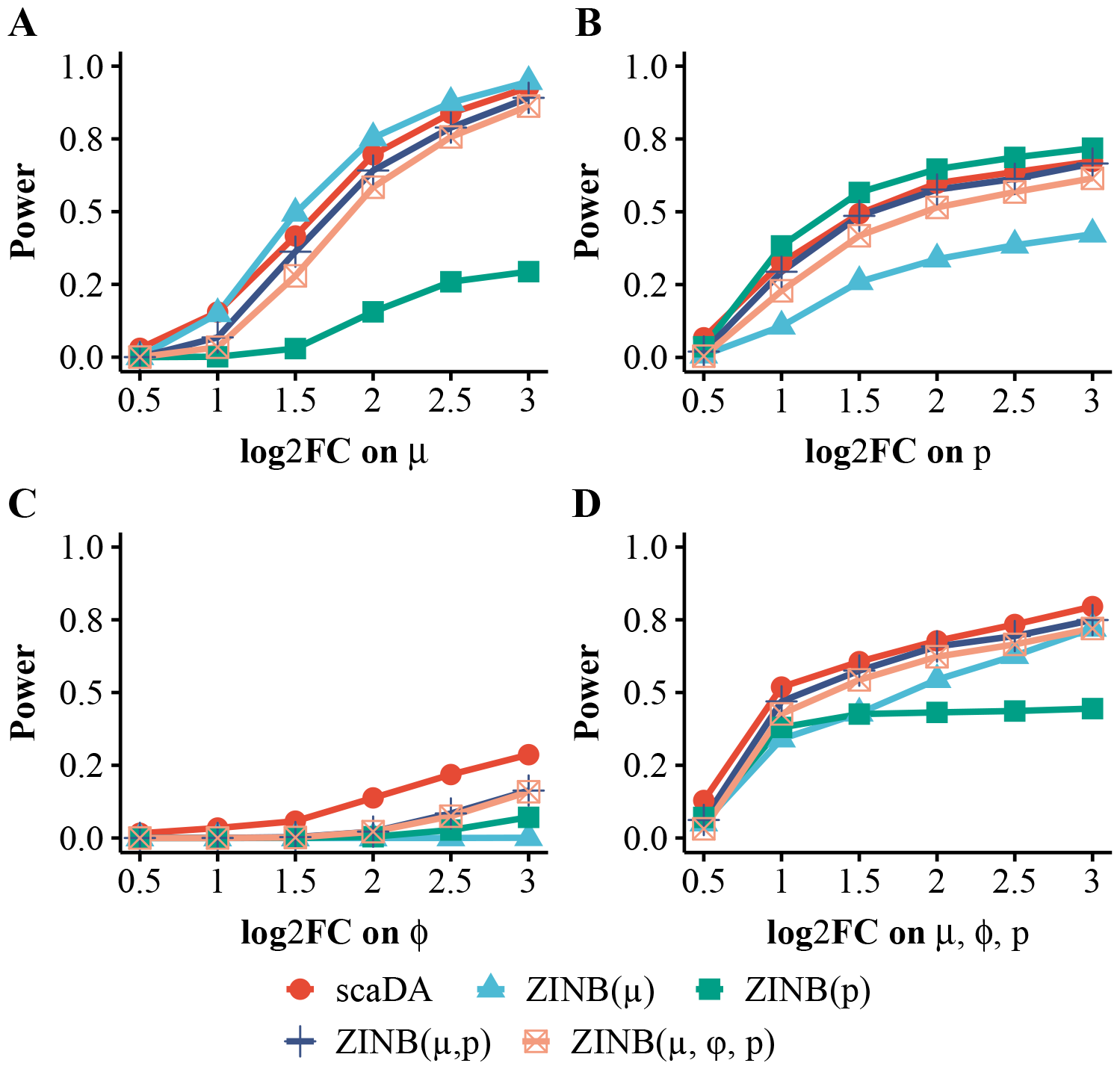
Power analysis for scaDA and other ZINB-based likelihood ratio tests. Baseline values of three parameters are estimated in the cell type “Granule neuron” from “Human Brain 3K”. For differential peaks, we fix the parameter values in Group 1 and vary the parameter values in Group 2 with different effect sizes of selected parameters to create power curves. To create parameter values in Group 2, we multiply and divide an effect size in terms of log2 fold change from the baseline values for differential peaks in equal proportion. The log2 fold change of selected parameters is changed from 0.5 to 3.0 with a step size of 0.5. Using the parameter setting in the four scenarios, we assume 20% differential peaks and simulate the read counts based on ZINB for 4000 peaks across 100 cells in each group. **A**. Scenario 1: only mean difference between two groups **B**. Scenario 2: only prevalence between two groups. **C**. Scenario 3: only dispersion difference between two groups. **D**. Scenario 4: difference of all three parameters between two groups.

When the underlying differential accessibility between the two groups is only driven by the mean difference (Scenario 1, Fig 3A) or prevalence difference (Scenario 2, Fig 3B), the corresponding tests, which include ZINB(*μ*) only testing on mean difference and ZINB(*p*) only testing on prevalence difference, are most powerful. Interestingly, for both Scenario 1 & 2, ZINB(*μ, p*) outperforms ZINB(*μ, ϕ, p*). Though both composite tests involve the parameters that drive the underlying differential accessibility between two groups, i.e. mean difference for Scenario 1 and prevalence difference for Scenario 2, ZINB(*μ, p*) has one less degree freedom and therefore is more powerful, especially considering *ϕ* in the composite test does not contribute to the underlying differential accessibility in Scenario 1 & 2. Notably, scaDA performs second best in both Scenario 1 & 2, which achieves slightly lower power than ZINB(*μ*) (Fig 3A) and ZINB(*p*) (Fig 3B). Notably, scaDA performs better than both ZINB(*μ, p*) and ZINB(*μ, ϕ, p*). Though both scaDA and ZINB(*μ, ϕ, p*) adopt the same composite hypothesis test with 3 d.f., scaDA is more powerful by benefiting from both dispersion shrinkage and refined estimates of mean and prevalence. Importantly, the performance of scaDA is not compromised by adding one more degree of freedom compared to ZINB(*μ, p*). Not surprisingly, ZINB(*p*), which tests on prevalence difference, performs worst in Scenario 1, where differential accessibility between the two groups is only driven by mean difference. Similarly, ZINB(*μ*), which which tests on mean difference, has the worst performance in Scenario 2, where differential accessibility between the two groups is only driven by prevalence difference. Nevertheless, ZINB(*μ*) still has power and the power increases when the effect size of prevalence difference increases. Similarly, ZINB(*p*) also obtains power and the power increases when the effect size of mean difference increases. This observation indicates a possible correlation between *μ* and *p*. Indeed, the prevalence is close to the mean for scATAC-seq data with sparse and low read count. If scATAC-seq data is binarized, the prevalence is equivalent to the mean.

In Scenario 3 (Fig 3C), it is not surprising that scaDA, which is the only test on dispersion difference, is the most powerful and holds a clear advantage over other methods when underlying differential accessibility between the two groups is only driven by dispersion difference. This observation indicates the necessity to include the dispersion parameter in the composite hypothesis test when the dispersion difference exists. The existence of dispersion difference among cell types is further validated by the real data exploration in “Human Brain 3K” (Fig 2C). Notably, even though both scaDA and ZINB(*μ, ϕ, p*) have 3 d.f., scaDA improves the power significantly compared to its counterpart. Moreover, it is also unsurprising to observe ZINB(*μ, p*) and ZINB(*μ, ϕ, p*), which do not involve dispersion parameter in the test, perform worst. In Scenario 4 (Fig 3D), when the underlying differential accessibility between the two groups is driven by the difference of all three parameters, scaDA achieves the highest power across all effect sizes consistently. Not surprisingly, the second best-performed methods are ZINB(*μ, p*) and ZINB(*μ, ϕ, p*), which are both composite tests involving multiple parameters simultaneously. Simple hypothesis tests on only one parameter such as ZINB(*μ*) and ZINB(*p*), are inferior to the composite tests such as ZINB(*μ, p*) and ZINB(*μ, ϕ, p*).

Overall, it is evident that scaDA exhibits the most robust performance across different scenarios and this is especially important in real data analysis because the factors that drive underlying differential accessibility between the two cell types are unknown. We further increase the sample size in each group to 200 and 300 respectively and repeat the simulations to evaluate the performance of scaDA and other methods under different sample sizes (Supplementary **Figure S2, Figure S3**). As expected, the power of all methods increases and the difference of method performance diminishes when the sample size increases. However, scaDA still remains the top-performed method and the advantage is more evident when dispersion is involved in the composite hypothesis testing.

### Power analysis for scaDA and published methods

We compare scaDA to state-of-the-art published approaches in the same simulation strategy as aforementioned. These approaches include methods for DE analysis of bulk RNA-seq such as NegBin and edgeR, methods for DE analysis of scRNA-seq such as MAST, and methods for DA analysis of scATAC-seq such as scATAC pro and Signac. Overall, scaDA is the most powerful test across all effect sizes in the four scenarios, with the only exception of Scenario 2, where it ranks second to scATAC-pro. Interestingly, MAST, which is a method developed for DE analysis of scRNA-seq data, performs second to scaDA in most scenarios (Fig 4). Specifically, in Scenario 1 (Fig 4A), scaDA ranks top, followed by Signac, edgeR, and NegBin. MAST and scATAC-pro are less powerful compared to other methods. In Scenario 2 (Fig 4B), scaDA is only slightly less powerful than scATAC-pro, closely followed by MAST. NegBin, edgeR, and Signac have an overall poor performance. In Scenario 3 (Fig 4C), scaDA has a clear advantage over other methods by achieving significantly higher power. Except for MAST, other methods lack statistical power to detect differential peaks. This observation further confirms the importance of including the dispersion parameter in the composite hypothesis test if underlying differential accessibility is driven by dispersion difference. In Scenario 4 (Fig 4D), scaDA still markedly outperforms other methods by achieving the highest power, followed by MAST, while other methods have comparable performance.

**Figure 4:**
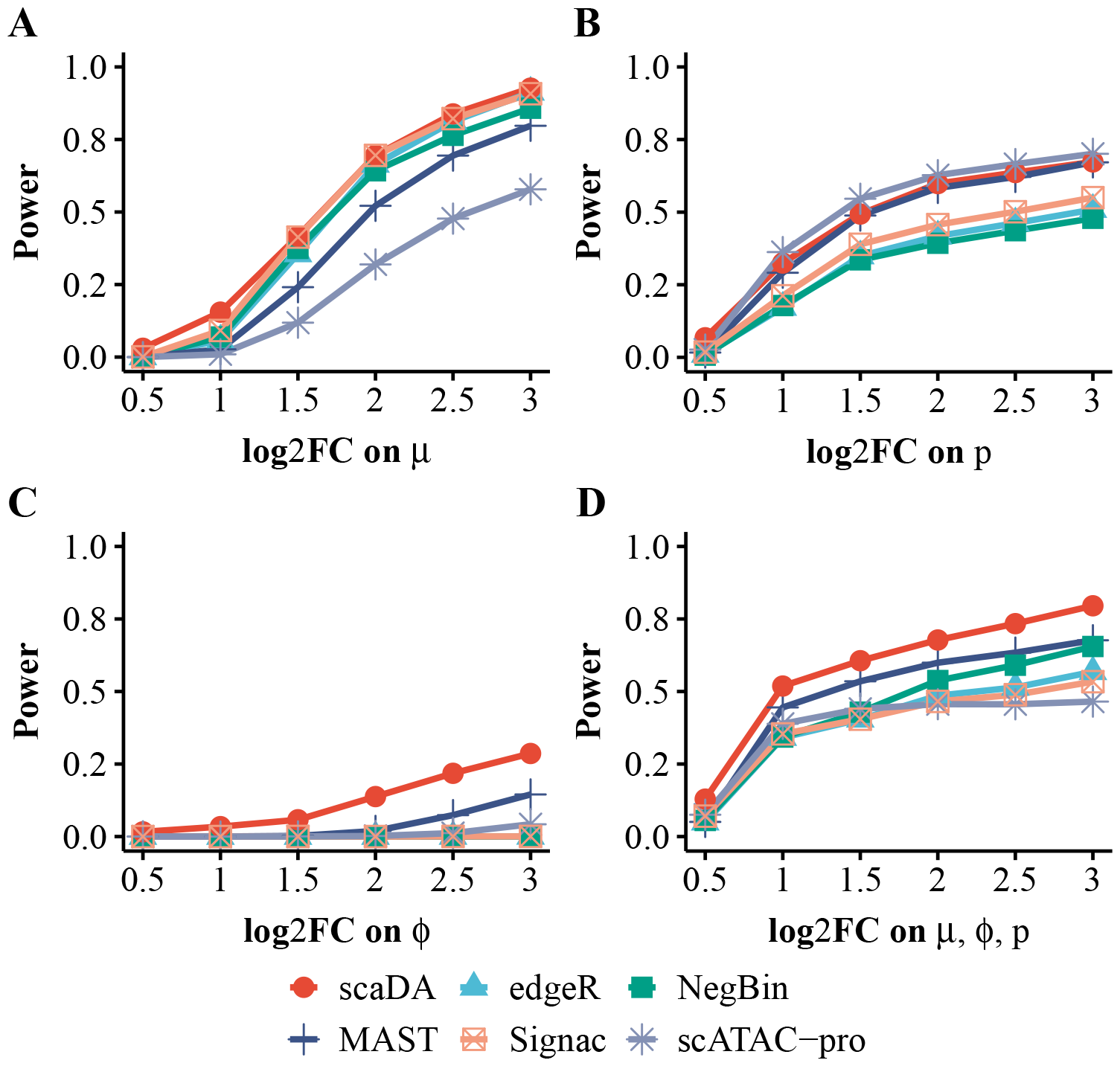
Power analysis for scaDA and published methods. Baseline values of three parameters are estimated in the cell type “Granule neuron” from “Human Brain 3K”. For differential peaks, we fix the parameter values in Group 1 and vary the parameter values in Group 2 with different effect sizes of selected parameters to create power curves. To create parameter values in Group 2, we multiply and divide an effect size in terms of log2 fold change from the baseline values for differential peaks in equal proportion. The log2 fold change of selected parameters is changed from 0.5 to 3.0 with a step size of 0.5. Using the parameter setting in the four scenarios, we assume 20% differential peaks and simulate the read counts based on ZINB for 4000 peaks across 100 cells in each group. **A**. Scenario 1: only mean difference between two groups **B**. Scenario 2: only prevalence between two groups. **C**. Scenario 3: only dispersion difference between two groups. **D**. Scenario 4: difference of all three parameters between two groups.

To evaluate the performance of all methods under different sample sizes, we repeat the simulations by increasing the sample size in each group to 200 and 300 respectively (Supplementary **Figure S4** and **Figure S5**). The power of all methods increases and the performance of all methods tends to converge when the sample size increases. Nevertheless, we observe a consistent trend that scaDA is superior to other methods and the advantage is more evident when the dispersion parameter is considered in the composite hypothesis testing.

### FDR control analysis

Besides the power analysis, we compare scaDA to both ZINB-based LRT tests and published methods in terms of FDR control. This is done by plotting the observed FDR against the nominal FDR level (Fig 5). Without loss of generality, we perform the evaluation in Scenario 4, which is a general case, and choose 2.5 as the log2 fold change for all three parameters. Compared to ZINB-based LRT tests, scaDA obtains the most accurate FDR estimation. The observed FDR of ZINB(*μ*) and ZINB(*p*) is overall slightly descended relative to the nominal level, indicating the test is slightly conservative. Notably, ZINB(*μ, ϕ, p*) and ZINB(*μ, p*) are overly conservative and give pessimistic results. Compared to published methods, scaDA has the top performance by controlling the observed FDR closest to the nominal level. All other methods such as scATAC-pro, edgeR, and MAST has an overly conservative FDR estimation. Notably, NegBin exhibits substantial inflated FDR, and the FDR curve exceeds the plotting range as the nominal FDR level increases (Fig 5B).

**Figure 5:**
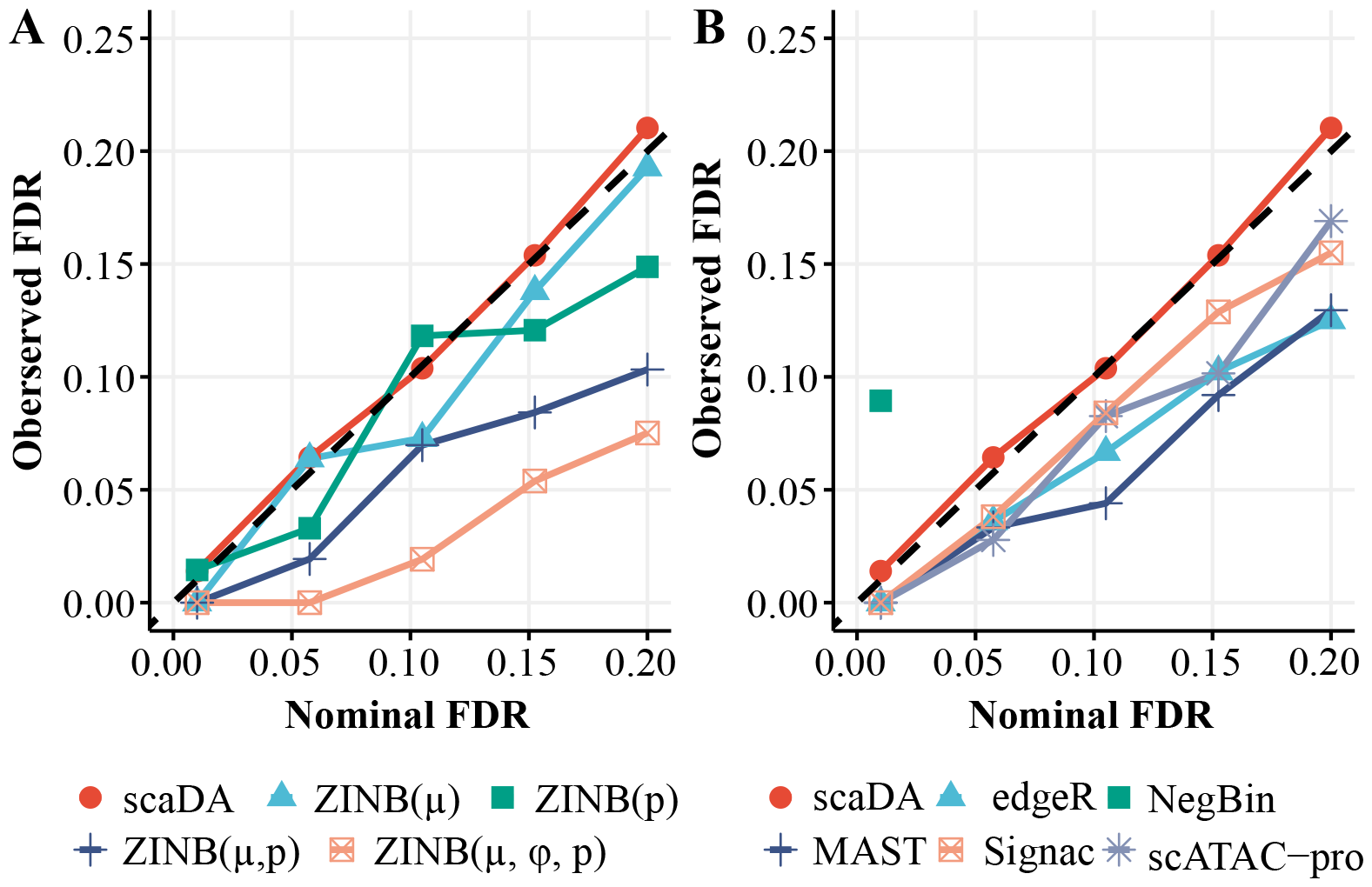
FDR control analysis. scaDA is compared to both ZINB-based LRT tests and published methods. Using the same simulation strategy in Scenario 4 (log2FC=2.5), we assume 20% differential peaks and simulate the read counts based on ZINB for 4000 peaks across 100 cells in each group. The observed FDR is plotted against the nominal FDR level. **A**. scaDA is compared to ZINB-based LRT tests. **B**. scaDA is compared to published methods.

Similarly, we perform the FDR control analysis when the sample size in each group increases to 200 and 300 respectively (Supplementary **Figure S6** and **Figure S7**). Compared to ZINB-based LRT tests, scaDA still achieves the best FDR control and ZINB(*μ*) becomes more conservative in FDR estimation (Supplementary **Figure S6**(A) and **Figure S7**(A)). Compared to published methods, scaDA is still the top-performed method though the performance of all methods tends to converge when the sample size increases. Notably, NegBin and edgeR, which are two methods developed for DE analysis of bulk RNA-seq, suffer a substantially inflated FDR, and the FDR curve exceeds the plotting range as the nominal FDR level increases (Supplementary **Figure S6**(B) and **Figure S7**(B)). Overall, all the evidence suggests that scaDA performs the best FDR control.

### Parameter estimation

We notice that scaDA outperforms its closest counterpart ZINB(*μ, ϕ, p*) in all simulation scenarios for both improved power and better FDR control even both composite hypothesis tests focus on the same set of parameters mean, dispersion, and prevalence. The key difference between scaDA and ZINB(*μ, ϕ, p*) lies in dispersion shrinkage and iterative refinement of mean and prevalence, which may lead to a better parameter estimation that is responsible for improved performance in both power analysis and FDR control.

To validate this, we evaluate the accuracy of parameter estimation for both methods. The correlation between the true dispersion (*ϕ*) and the estimated dispersion 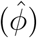 for both ZINB(*μ, ϕ, p*) and scaDA are shown in Fig 6A. Consequently, scaDA yields a better correlation by achieving a higher Pearson correlation coefficient of 0.443, which is higher than 0.303 obtained by ZINB(*μ, ϕ, p*). In addition, scaDA consistently obtains lower MSEs in all three parameters (Fig 6B), which indicates the effectiveness of parameter estimation by scaDA.

**Figure 6:**
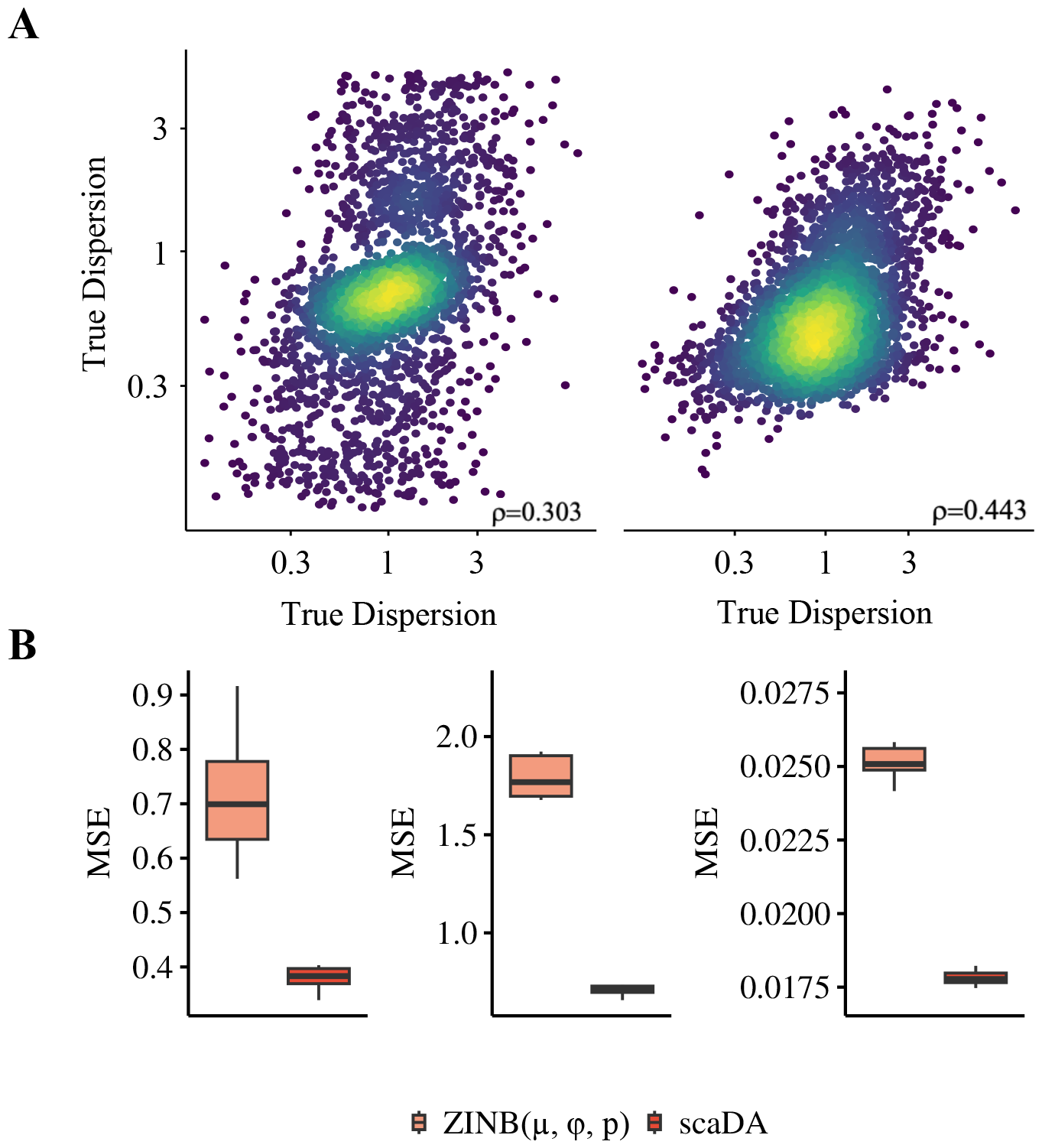
Parameter estimation. Using the same simulation strategy in Scenario 4 (log2FC=2.5), we assume 20% differential peaks and simulate the read counts based on ZINB for 4000 peaks across 100 cells in each group. **A**. True dispersion is plotted against the estimated dispersion from ZINB(*μ, ϕ, p*) and scaDA on a log scale plot. **B**. MSE calculated between true and estimated mean, dispersion, and prevalence from ZINB(*μ, ϕ, p*) and scaDA.

### Evaluating DA analysis using three sc-multiome data

#### Overview of evaluation strategy

We perform DA analysis of scaDA and published methods in three sc-multiome datasets. The analyses for “Human Brain 3K” and “Human PBMC 10K” focus on identifying differential peaks between one cell type and other cell types. Stratified sampling is adopted to ensure balanced sample sizes between the two groups. Detecting differential peaks in the first type of DA analysis is crucial for hypothesis generation, where cell-type specific peaks can be identified for cell type annotation. The analyses for “Human AD” focus on detecting the differential peaks between two disease stages (i.e., AD and control) of each cell type. The second type of DA analysis can be widely used to identify disease-associated regions, which can be used for disease prevention and treatment.

Since the gold standard for measuring differential peaks between cell types is not available, we evaluate the differential peaks by leveraging the status of nearby differential gene expression because chromatin accessibility near the promoters is positively correlated with gene expression. Based on the data type of sc-multiome data, we qualify the gene expression in two ways. For scmultiome data with ATAC and RNA profiled in the same cell, we use the profiled gene expression directly. For sc-multiome data with ATAC and RNA not profiled in the same cell, we estimate the gene expression by leveraging ATAC-seq counts in the 2 kb-upstream and downstream region of transcriptional start sites (TSS). Specifically, we use “GeneActivity()” function in the Signac package for the estimation of gene expression based on ATAC-seq count. Once the gene expression is quantified, we perform DE gene analysis using the Wilcoxon rank-sum test. We adopt the Wilcoxon test is because the non-parametric test is more conservative than its parametric counterparts, reducing the potential false positives. Consequently, genes with FDR less than 0.05 and the absolute value of log2 fold change in the first quantile are deemed as DE, and others are deemed as non-DE. Next, we keep all peaks that are in proximity to both DE and non-DE genes and the proximity is quantified as peaks within the window of 50kb with TSS in the center. Peaks the 50kb window of corresponding DE genes are considered as true differential and within the 50kb window of non-DE genes are deemed as non-differential. Finally, we perform DA analysis using scaDA and published methods. Peaks with FDR less than 0.05 and an absolute value of log2 fold change larger than 0.5 are deemed as differential peaks and otherwise non-differential.

As the goal of DA analysis is to have as many true positives as possible among the top-ranked differential peaks, we adopt the proportion of true differential peaks among top-ranked differential peaks, named true discovery rate (TDR), which has been widely used for quantifying the differential biological signals [21, 16], as the performance measure in the real data analysis. Specifically, we calculate TDR in the top-ranked peaks ranging from 20% to 100% with a step size of 20%. When all peaks are used i.e., 100%, TDR is equivalent to the True Positive Rate (TPR), which is also the statistical power.

#### Human Brain 3K

We employ the Seurat 10x sc-multiome data analysis pipeline to process “Human Brain 3K”, which profiles DNA accessibility and gene expression in the same cells, by performing quality control, data transformation, dimension reduction, and cell type annotation. For the quality control, we keep cells with counts of ATAC reads less than 100k but larger than 1k and counts of RNA reads less than 25k but larger than 1k. In addition, we keep cells with nucleosome signal scores less than 2, TSS enrichment scores larger than 1 and percent of mitochondrial gene expression less than 20%. For data transformation, sctransform [22] is applied to RNA modality and TF-IDF is used for ATAC modality. For dimension reduction, PCA and SVD are performed for RNA and ATAC modality respectively. Finally, Weighted Nearest Neighbor (WNN) analysis is performed for UMAP visualization and clustering by integrating both RNA and ATAC-seq modalities. Consequently, 14 distinct clusters have been identified with number of cells and cell type proportions reported in Supplementary **Table S1**. For cell type annotation, DE analysis is performed between each cluster and all the other clusters. Top DE genes for each cluster are identified and compared to reported gene markers from a published study on human cerebellar development [20].

As aforementioned, DE analysis is performed using RNA modality and true differential peaks are defined as peaks proximate to DE genes. TDR is calculated by comparing the differential peaks identified by each DA method and true differential peaks. Consequently, we plot the TDR across cell types at different levels of top-ranked differential peaks identified by each method (Fig 7). Overall, all methods exhibit the highest mTDR (mean TDR) across cell types in top 20% peaks and experience declined mTDR as the top percentage increases, which indicates that all methods are more powerful to identify true differential peaks in top-ranked peaks. Importantly, scaDA achieves the best performance by obtaining the highest mTDR in top 20% differential peaks (0.81 for scaDA, 0.72 for Signac, 0.72 for scATAC pro, 0.67 for MAST, 0.64 for NegBin, 0.67 for edgeR). The trend consistently holds across all levels of top percentages. When the top percentage equals 100%, TDR is equivalent to TPR (i.e. power). It is clear that scaDA is most powerful (0.51 for scaDA, 0.34 for Signac, 0.34 for scATAC pro, 0.33 for MAST, 0.22 for NegBin, 0.28 for edgeR). In addition, TDR demonstrates a large variability across cell types at different levels of top percentages. However, scaDA still holds less variability than majority of compared methods except for NegBin in top 20% peaks (0.03 for scaDA, 0.09 for Signac, 0.08 for scATAC pro, 0.10 for MAST, 0.05 for NegBin, 0.08 for edgeR). Signac, scATAC-pro, MAST and edgeR generally yield comparable performances but exhibit a higher variability than scaDA and NegBin. NegBin obtains the second lowest mTDR but lowest variability across all levels of top percentages. Interestingly, NegBin has less variability than its closest counterpart edgeR. The summary of mean and variability of TDR of all methods across all levels of top percentages can be found in Supplementary **Table S2, Table S3**. Moreover, we evaluate the power of all methods for each cell type (Fig 7B). scaDA performs the best by obtaining the highest power in 6 out of 8 cell types, which is followed by Signac with highest power in 2 out of 8 cell types (Supplementary **Table S4**). We further explore the TDR of all methods for each cell type at different levels of top percentages and find scaDA consistently have overall best performance (Supplementary **Figure S8**)

**Figure 7:**
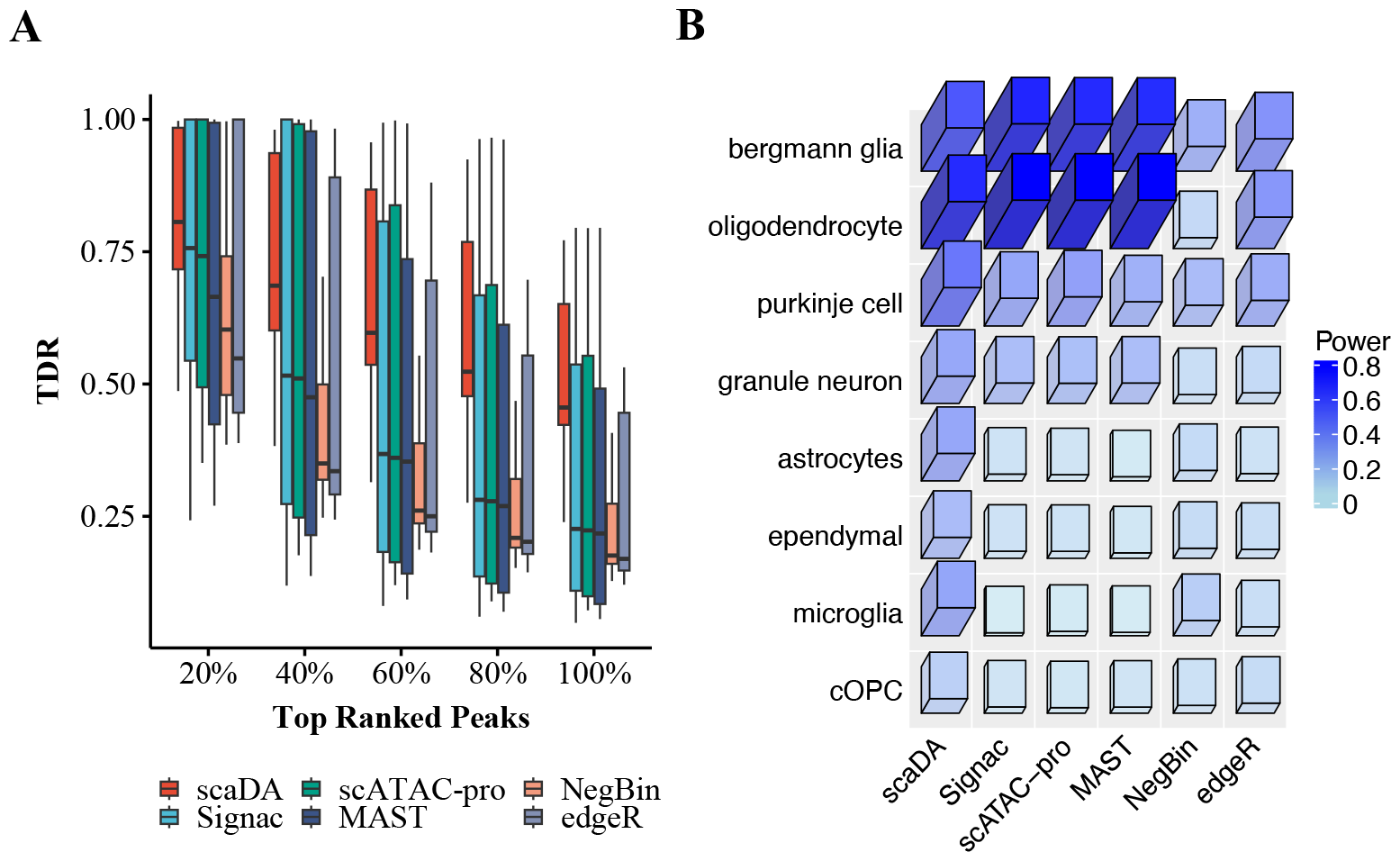
Human Brain 3K. **A**. True Discovery Rate (TDR) is reported across 8 cell types for all methods at different levels of top percentages (i.e., 20%, 40%, 60%, 80%, and 100%) **B**. Power (TDR at 100%) across 8 cell types for all methods.

#### Human PBMC 10K

Similarly, Seurat 10x sc-multiome data analysis pipeline is used to process “Human PBMC 10K”, which profiles DNA accessibility and gene expression in the same cells, by performing quality control, data transformation, dimension reduction, and cell type annotation. In particular, cell type annotation is performed using PBMC reference scRNA-seq data [23]. As a result, 20 clusters are identified and annotated, and cell type with cell number less than 100 are excluded resulting 14 cell types for downstream analysis. Cell type proportion and number of cells in each cell type are reported in Supplementary **Table S5**.

Similar to before, true differential peaks are defined as peaks proximate to DE genes. TDR is calculated by comparing the differential peaks identified by each DA method and true differential peaks. As a result, TDR across 14 cell types at different levels of top-ranked differential peaks, which are identified by each method, are demonstrated in Fig 8. It is expected that the performance of all methods decrease when the top percentage increases. Notably, scaDA obtains the highest mTDR in the top 20% differential peaks (0.57 for scaDA, 0.49 for NegBin, 0.45 for edgeR, 0.45 for Signac, 0.43 for scATAC pro and 0.40 for MAST). Moreover, scaDA achieves the highest power (0.24 for scaDA, 0.16 for NegBin, 0.14 for edgeR, 0.10 for Signac, 0.10 for scATAC pro and 0.09 for MAST). In addition, the trend persists across all levels of top percentages. Besides achieving the highest mTDR, scaDA also holds the smallest variability of TDR across cell types among all methods in top 20% differential peaks (0.02 for scaDA, 0.07 for NegBin, 0.06 for edgeR, 0.07 for Signac, 0.07 for scATAC pro and 0.08 for MAST). For scaDA, the trend of yielding the smallest variability of TDR across cell types is consistent across all levels of top percentages. In addition, the superiority of scaDA is more evident for top-ranked differential peaks (e.g., 20%, 40%). In contrast, other methods suffer a larger variability of TDR across cell types in top-ranked differential peaks (e.g., 20%, 40%). The variability of other methods will decrease when the percentage increases. The summary of the mean and variability of TDR of all methods across all levels of top percentages can be found in Supplementary **Table S6, Table S7**. We further investigate the performance of all methods for each cell type. scaDA obtaining the highest power in 9 out of 14 cell types and second highest in 5 out of 14 cell types (Fig 8B, Supplementary **Table S8**). In addition, the superiority of scaDA persists at different levels of top percentage (Supplementary **Figure S9**).

**Figure 8:**
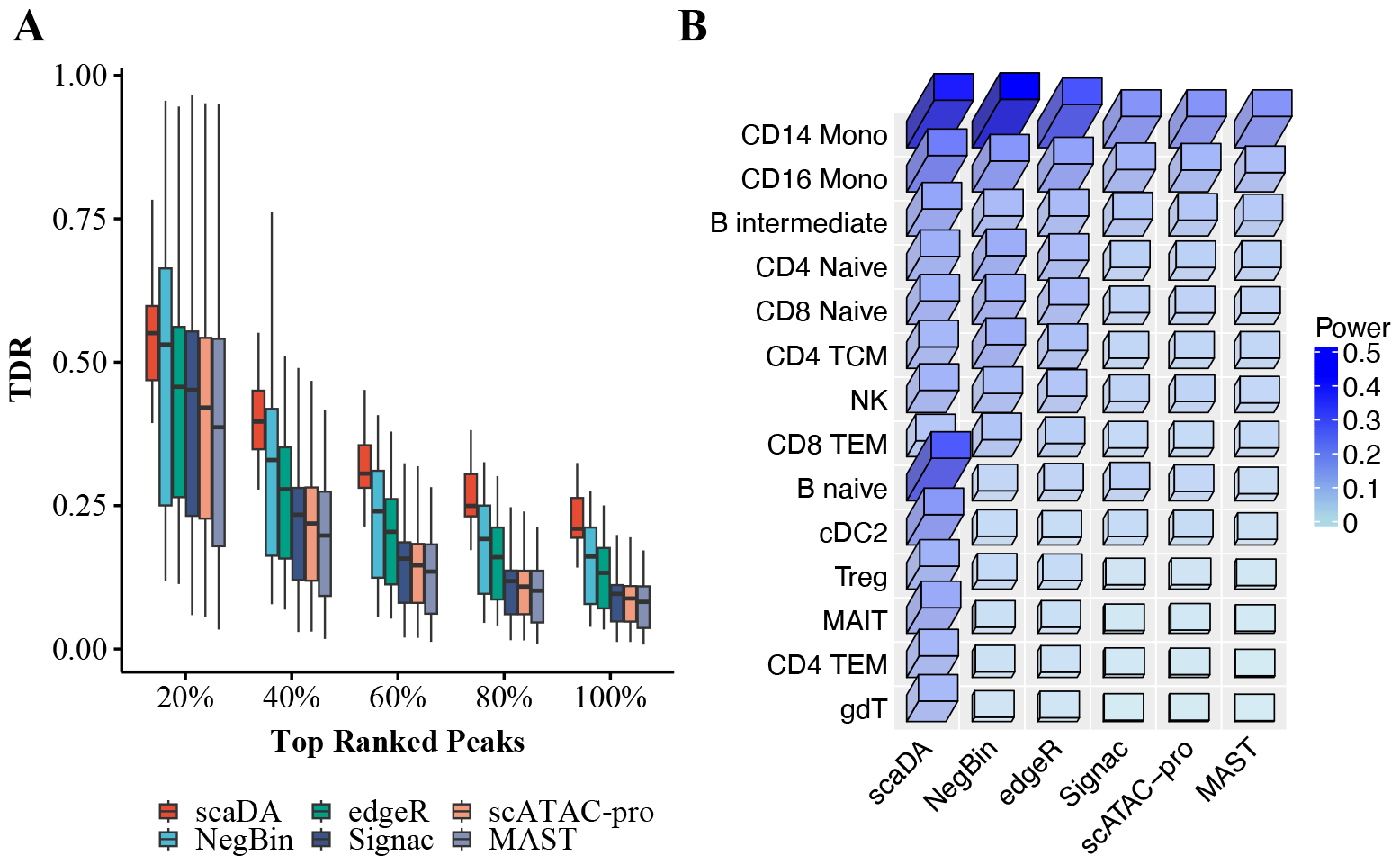
Human PBMC 10K. **A**. True Discovery Rate (TDR) across 14 cell types for all methods at different levels of top percentages (i.e., 20%, 40%, 60%, 80%, and 100%) **B**. Power (TDR at 100%) across 14 cell types for all methods.

#### Human AD

“Human AD” consists of 12 AD and 8 healthy control samples across 3 batches. Cell type annotation has been provided for all cells in the metadata. Seven major cell types are provided, which include Astrocytes (ASC), Excitatory neurons (EX), Inhibitory neuron (INH), Microglia (MG), Oligodendrocytes (ODC), Olig. progenitors (OPC) Pericytes/endothelial (PER.END). Due to few cells in PER.END, we remove PER.END from the following analysis. As the RNA and ATAC modality are profiled in separate scATAC-seq and scRNA-seq data, we use the ATAC modality to infer “pseudo” gene expression for DE analysis to prevent the potential domain shift of data modality. We employ the Signac 10x scATAC-seq data analysis pipeline to process the data by performing quality control, data transformation, dimension reduction, and cell type annotation. To reduce the batch effect, we perform cell type-specific DA analysis for each pair of one AD and one control sample within each batch. TDR is calculated as aforementioned accordingly.

Consequently, TDR across all cell types and pairwise comparison at different levels of topranked differential peaks are presented in Fig 9A. scaDA obtains the highest mTDR in the top 20% differential peaks (0.77 for scaDA, 0.65 for scATAC pro, 0.62 for MAST, 0.51 for edgeR, 0.45 for Signac, and 0.16 for NegBin). Similarly, scaDA performs much better than other methods by obtaining a significantly higher power (0.69 for scaDA, 0.34 for scATAC pro, 0.33 for MAST, 0.35 for edgeR, 0.21 for Signac, and 0.09 for NegBin). A persistent trend, which scaDA consistently obtains the highest mTDR, is observed for across all levels of top percentages. Except for NegBin, which has the worst performance in terms of mTDR, scaDA achieves the smallest variability of TDR across all cell types and pair comparison among all methods in top 20% differential peaks (0.10 for scaDA, 0.17 for scATAC pro, 0.17 for MAST, 0.16 for edgeR, 0.19 for Signac and 0.07 for NegBin). The trend of variability of TDR across all cell types and pair comparison is consistent at all levels of top percentages, where scaDA and NegBin have a small variability but other methods suffer a large variability. The summary of the mean and variability of TDR across all cell types and pair comparison at different levels of top percentages can be found in Supplementary **Table S10, Table S11**. In addition, for each method, the mean power of all pairwise comparison is reported for each cell type (Fig 9B). Remarkably, scaDA is most powerful by obtaining the highest power in 6 out of 6 cell types. Following scaDA, scATAC-pro achieves second highest power in 3 out of 6 cell types (Supplementary **Table S12**). For each method, mTDR of all pairwise comparison for each cell type at different levels of top percentages is explored (Supplementary **Figure S10**). Again, scaDA achieves the best performance by obtaining an overall highest cell type-specific mTDR.

**Figure 9:**
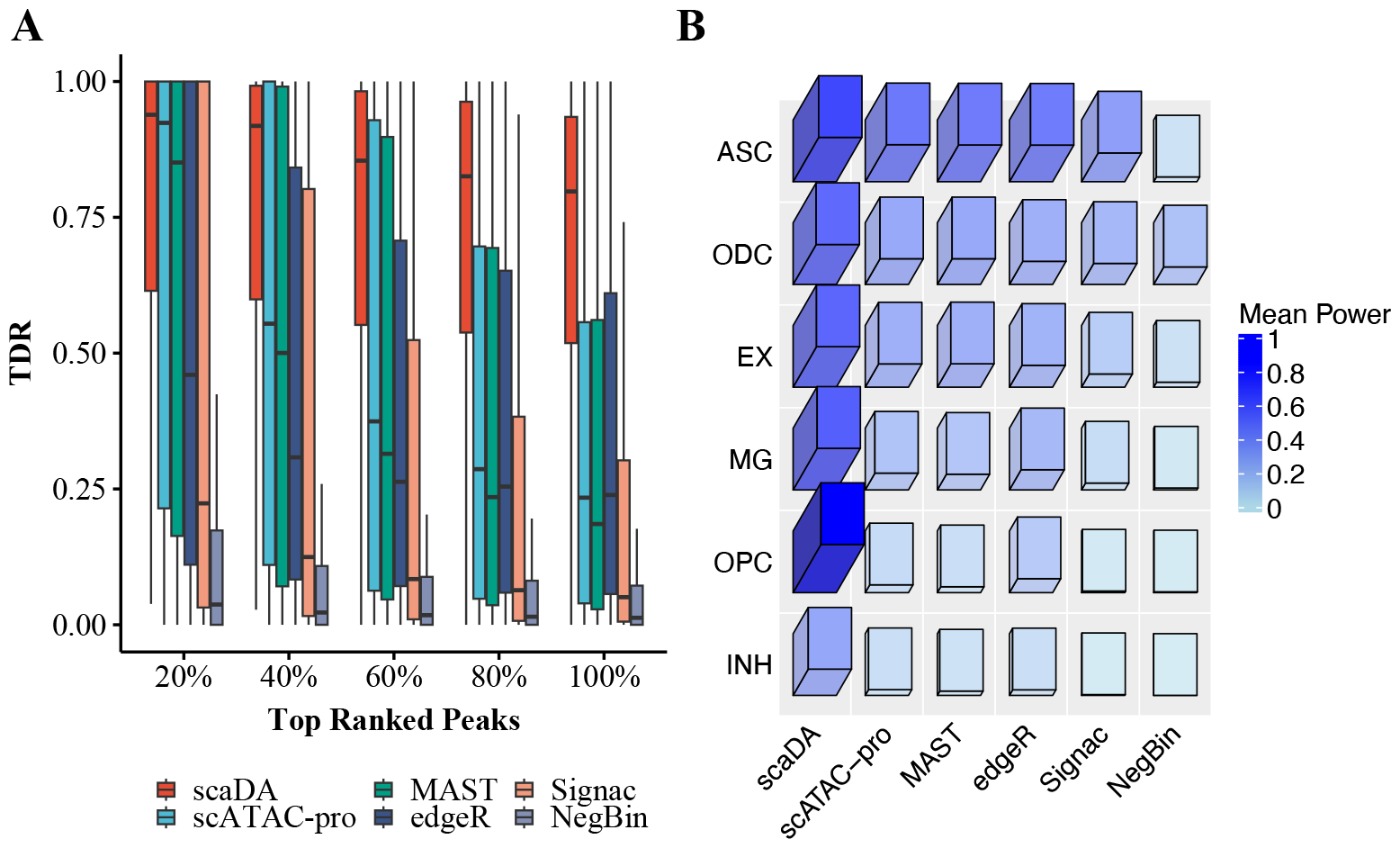
Human AD. **A**. Mean True Discovery Rate (TDR) across all 6 cell types and pairwise comparison for all methods at different levels of top percentages (i.e., 20%, 40%, 60%, 80%, and 100%) **B**. Mean Power (TDR at 100%) across all 6 cell types and pairwise comparison for all methods.

### Gene Ontology analysis

We perform the Gene ontology (GO) analysis for differential peaks identified by each method (FDR*<*0.05, abs(log2FC)*>*0.5). For the GO analysis, we choose microglia from “Human AD” for DA analysis. Microglia are innate immune cells of the myeloid lineage that reside in the central nervous system (CNS) and are key cellular mediators of neuroinflammation. It is well-known that microglia has demonstrated a key role in the pathogenesis of AD by generating adaptive immune response [24]. Therefore, a top-performed method is expected to identify differential peaks that are more functionally enriched in neural and immune-related GO terms.

First, we assign identified differential peaks to protein-coding genes, which will be used for GO analysis. The selection of the genes is based on the positional overlap between the differential peaks and 50kb extended region of TSS with TSS in the middle. Next, we perform GO term analysis using the “enrichGO” function in “clusterProfiler” R/Bioconductor package [25] with default parameter setting such as pAdjustMethod = “BH” and qvalueCutoff = 0.05. We compare scaDA to the published methods including NegBin, edgeR, MAST, scATAC-pro and Signac. For each method, we perform the DA analysis between one AD and one control sample by pairwise comparison within each of the three batches, which results in 34 sets of differential peaks and 34 sets of GO analysis results. For each GO term, we employ Stouffer ‘s Z-score method [26], which combines p-values from 34 sets into a single p-value, and then apply multiple testing correlation to obtain FDR.

Consequently, the performance of all methods has been demonstrated by enrichment in 10 significant GO terms (FDR*<*0.05) (Fig 10), which include “cognition” and “learning or memory” related to clinical phenotype of AD involves progressive cognitive decline and memory loss [27]; “regulation of trans-synapic signaling” and “moderation of chemical synaptic transmission” related to synaptic transmission that promotes neurodegeneration [28]; “regulation of amyloid-beta formation” and “amyloid-beta metabolic process” related to Amyloid beta, which is one protein crucial to AD development [29]; “immunological memory process” and “negative regulation of adaptive immune response” related to microglia ‘s role in the pathogenesis of AD by generating adaptive immune response [24].

**Figure 10:**
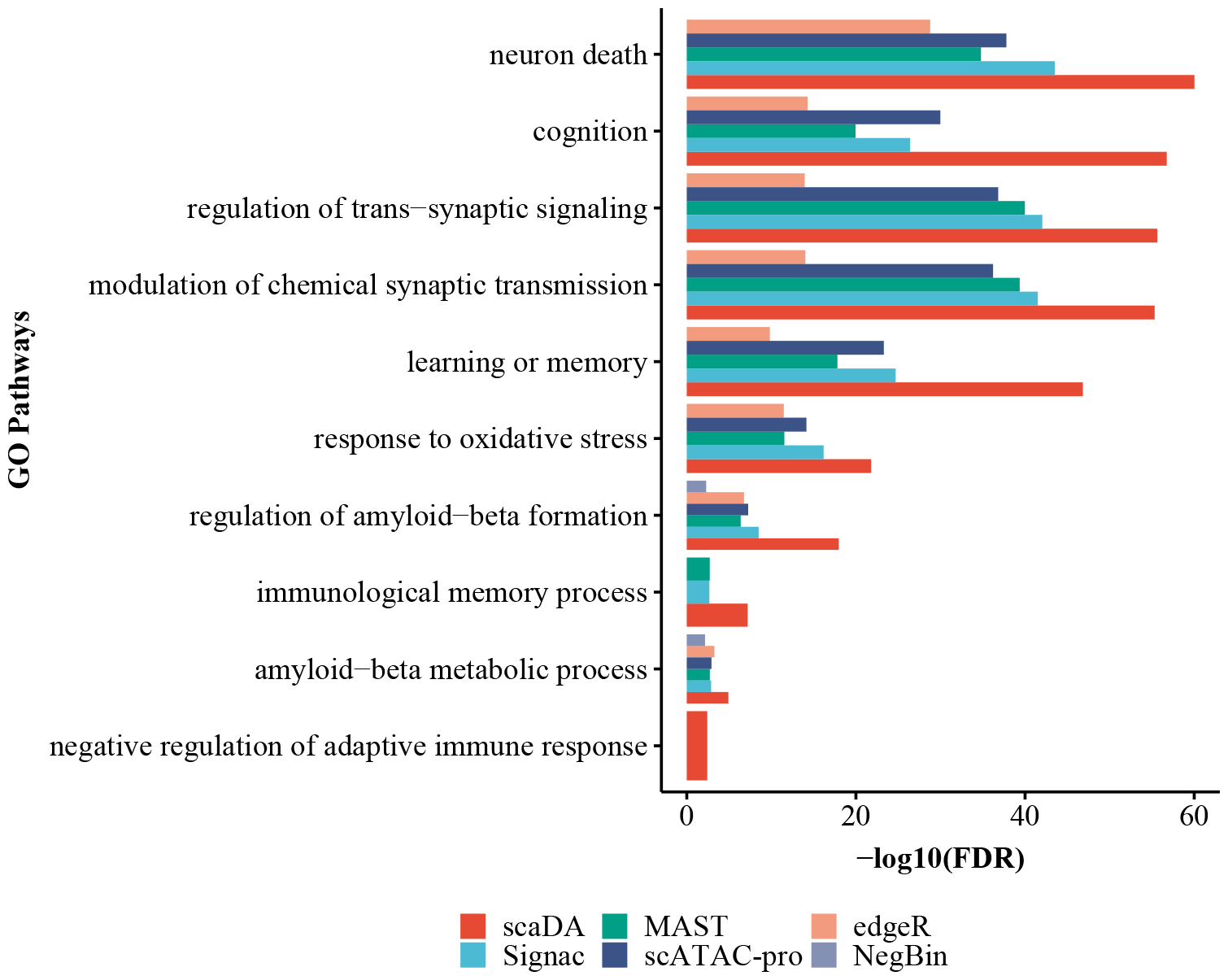
GO analysis for AD microglia. Gene ontology (GO) analysis is performed on differential peaks identified by scaDA and published methods. Microglia from “Human AD” is selected for the GO analysis.

The majority of all methods are enriched in the 10 key GO terms related to AD neurodegeneration and clinical phenotype of AD. Remarkably, scaDA stands out by obtaining the highest enrichment. Following scaDA, scATAC-pro, MAST and Signac, three methods developed for singlecell genomics, rank in the second tie. edgeR and NegBin, two methods developed for differential analysis of bulk RNA-seq, perform worst. Particularly, NegBin fails to identify differential peaks enriched in any of the GO terms. All the evidence demonstrates that scaDA is capable of identifying differential peaks enriched in GO terms related to neurogenesis and clinical phenotype of AD, which further validates scaDA is still the top-performed method compared to its counterparts by GO analysis.

### GWAS enrichment analysis

GWAS enrichment analysis aims to evaluate the enrichment of diseases/traits-associated GWAS SNPs in cell type-specific chromatin accessible regions. The analysis can help identify disease/trait-associated cell types and provide functional annotation and interpretation for GWAS SNPs [6].

Recently, GWAS enrichment analysis has been widely performed in several AD studies to evaluate the enrichment of AD-associated GWAS SNPs in chromatin accessible regions across major brain cell types [19, 30]. Enlightened by the previous studies, we adopt GWAS enrichment analysis to evaluate the differential peaks identified by all methods.

Similar to the GO analysis, we select microglia from “Human AD” for DA analysis and identify differential peaks for each method. For AD-associated GWAS SNPs, we collect AD-associated GWAS summary statistics in a study named “genome-wide association study by proxy (GWAX)” [31]. We identify 1302 positive AD-associated GWAS SNPs (p-value¡1×10^*−*4^) and generate negative control SNPs 10 times more than the positive set using the strategy from previous work [32, 33, 34]. Next, for each method, we count how many positive/negative SNPs within differential peaks/nondifferential peaks and construct a 2 by 2 contingency table. We then perform Fisher ‘s exact test to calculate the odd ratio (OR), confidence interval (CI) and p-value for the table. Similar to the GO analysis, for each method, we perform the DA analysis between one AD and one control sample by pairwise comparison within each of the three batches, which results in 34 sets of differential peaks and 34 sets of OR, CI and p-value. To integrate 34 sets of results and resolve the potential discrepancy, we perform a meta-analysis by adopting the R package “meta”. Consequently, we obtain one set of OR, CI and p-value for each method (Fig 11). Consequently, scaDA achieves the highest OR (logOR=1.97) and most significant finding (p-value=0.026). Interestingly, edgeR ranks second by achieving slightly lower OR and less significant pvalue (logOR=1.88 and pvalue=0.037). Though positive AD-associated GWAS SNPs are enriched in differential peaks identified by MAST and scATAC-pro (logOR=1.56 and 1.15), the finding is not statistically significant (pvalue=0.081 and 0.198). Surprisingly, NegBin and Signac show unfavorable performance by failing to detect differential peaks hit by positive AD-associated GWAS SNPs.

**Figure 11:**
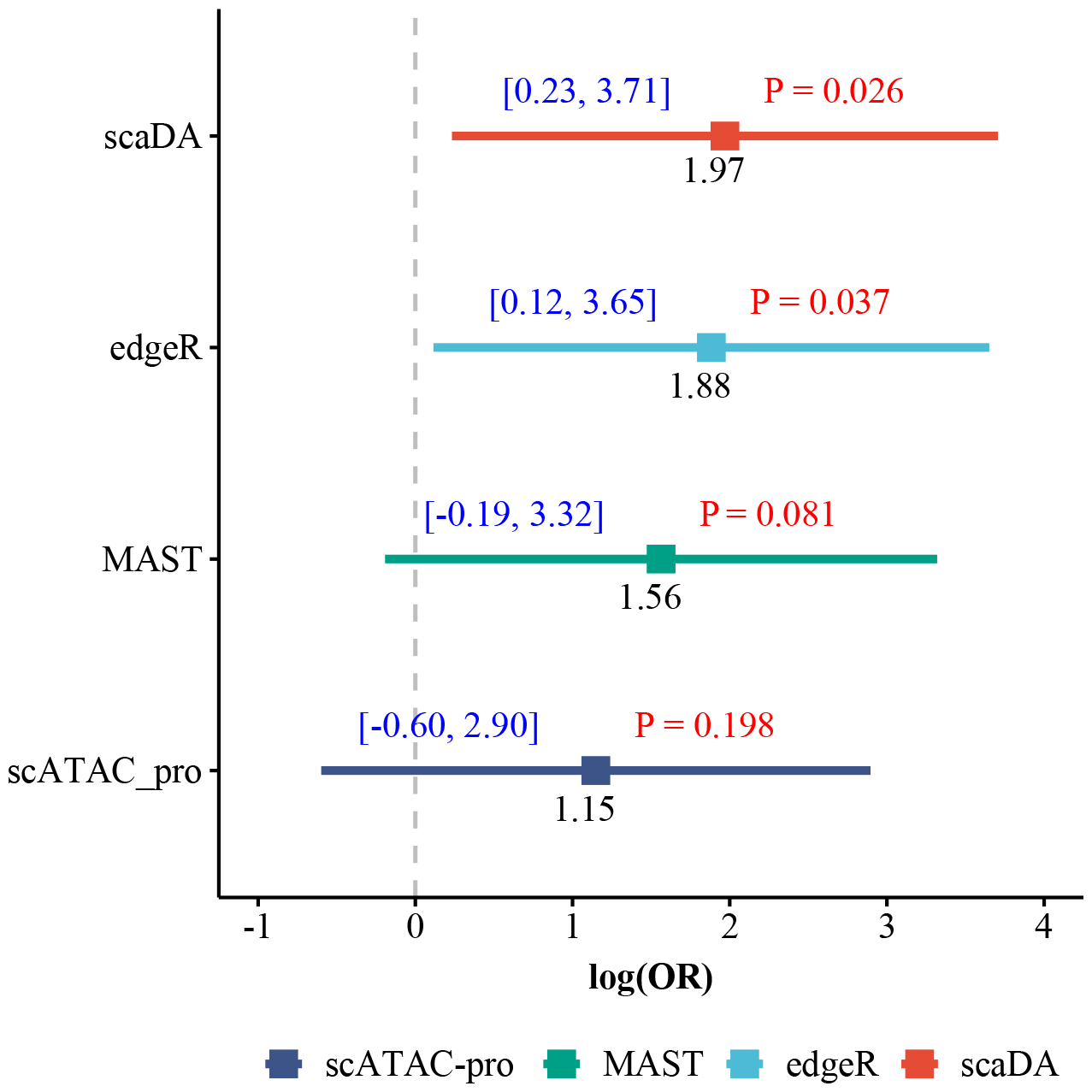
GWAS enrichment analysis. Microglia from “Human AD” is selected for DA analysis and identify differential peaks for each method. GWAS enrichment analysis is performed by evaluating the enrichment of AD-associated GWAS summary statistics in microglia-specific differential peaks. Results are presented in terms of logOR, CI and pvalue.

## Discussion

With the advent of scATAC-seq data, differential chromatin (DA) analysis is able to identify celltype specific chromatin accessible regions by comparing two cell types, or detect disease-associated chromatin accessible regions by the two-group analysis (e.g., disease and normal) for the same cell type. The DA regions can be further linked to dynamic cis-regulatory elements and regulated genes to have a comprehensive picture of gene regulatory activities. Statistical methods have been widely developed and used for performing DA analysis. Nonparametric methods, such as Wilcoxon rank-sum test, have been adopted by state-of-the-art scATAC-seq analysis pipelines such as Signac and scATAC-pro because of its robustness to distribution assumption. Nevertheless, Wilcoxon test is unable to adjust covariates and provide interpretable effect sizes. Other statistical methods such as MAST, which has been developed for differential expression analysis for scRNA-seq data, can be used directly for DA analysis. However, these methods may not be optimal for scATAC-seq data due to the different data characteristics such that scATAC-seq has more severe sparsity due to more peaks than genes and higher drop rates [11].

Considering the excessive zeros of scATAC-seq data, it is ideal to model the ATAC count data using a zero-inflated negative binomial model (ZINB). Motivated by real data exploration where we observe “distribution difference”, that is, distributions of all three parameters are different between multiple pairs of cell types, we decide to adopt composite hypothesis testing, which is more robust when the difference of chromatin accessibility is not determined by one single parameter. Collectively, we develop scaDA, a novel statistical framework based on ZINB model for DA analysis by joint considering mean, dispersion, and prevalence in a composite hypothesis testing framework. To improve the parameter estimation, scaDA incorporates a powerful empirical Bayes approach to derive a posterior estimate of dispersion, and further refines the estimate of mean and prevalence in an iterative fashion. To evaluate the effectiveness of scaDA, we perform a semi-parametric simulation based on one real scATAC-seq data, where mean and prevalence parameter values are randomly sampled from real data estimates and dispersion parameter values are generated from a log-normal distribution with hyper-parameters computed from the same real data. Importantly, we design four scenarios, where the differential chromatin accessibility is driven by each of the three parameters or three parameters at the same time.

Consequently, scaDA outperforms both ZINB-based LRTs and state-of-the-art published methods respectively. When comparing to ZINB-based LRTs that the differential chromatin accessibility is only driven by mean or prevalence, scaDA only slightly less powerful than ZINB(*μ*) and ZINB(*p*) respectively. However, when differential chromatin accessibility is driven by dispersion or three parameters, scaDA is most powerful. Importantly, scaDA outperforms its closest counterpart-ZINB model ZINB(*μ, ϕ, p*), which also performs composite test for three parameters. This observation can be explained by the additional benefit of improving the parameter estimation by dispersion shrinkage and parameter refinement for mean and prevalence, which is implemented by scaDA. Continuing the momentum, scaDA maintains top-performed method among state-of-the-art published methods, which ranges from DE methods for bulk-RNA and scRNA-seq to DA methods for scATAC-seq. Moreover, FDR control analysis confirms the superiority of scaDA by obtaining the best FDR control among both ZINB-based LRTs and published methods. We further explore different sample sizes for both power analysis and FDR control analysis. The overall trend persists that scaDA maintain the top performance. However, the advantage of scaDA diminishes as the benefit of increased sample size outweighs the gain from the more advanced method.

We apply scaDA to three sc-multiome data by leveraging the status of nearby differential expressed genes to evaluate differential peaks. We use true discovery rate (TDR), which is defined as proportion of true differential peaks among top-ranked differential peaks to evaluate the model performance. As a result, scaDA is most powerful by obtaining the highest mean and lowest variability of TDR across all cell types at different levels of top percentages. In addition, scaDA is the most frequently top-ranked method by power across all cell types. To evaluate the functional consequence of identified differential peaks, we perform Gene Ontology analysis on the differential peaks identified from microglia in “Human AD”. Consequently, scaDA successfully identifies differential peaks, which are most enriched in GO terms related to neurogenesis and clinical phenotype of AD. Furthermore, we perform GWAS enrichment analysis on identified differential peaks by leveraging a published AD-associated GWAS summary statistics. As a result, AD-associated GWAS SNPs are more enriched in differential peaks identified by scaDA than other published methods. Therefore, scaDA outperforms existing methods by both GO analysis and GWAS enrichment analysis, which further confirms the practical advantage of using scaDA in DA analysis.

Despite the favorable performance of scaDA, scaDA is limited in several aspects. Similar to existing methods, scaDA performs the DA analysis for each chromatin accessible region individually and independently. In fact, these peaks can be co-accessible, which can be predicted from scATACseq data using method such Cicero [35]. Thus, scaDA can be extended to test the differentially co-accessible peaks in a multivariate test. The multivariate test may help alleviate the multiple testing burden, increases the statistical power and provide novel biological findings. In addition, the current version of scaDA is limited to two-group comparison and our immediate plan is to extend scaDA to a multifactor design. This extension is crucial as the experimental design for scATAC-seq becomes sophisticated, where multiple batches, conditions, platforms and treatments can be considered simultaneously. We will explore these directions to extend and improve scaDA in the future work.

## Acknowledgments

This work was supported by the National Institute of General Medical Sciences of the National Institutes of Health [Award Number R35GM142701 to L.C.].

