## Supplementary materials for "scaDA: A Novel Statistical Method for Differential Analysis of Single-Cell Chromatin Accessibility Sequencing Data"

#### Supplemental Figures

##### Simulation

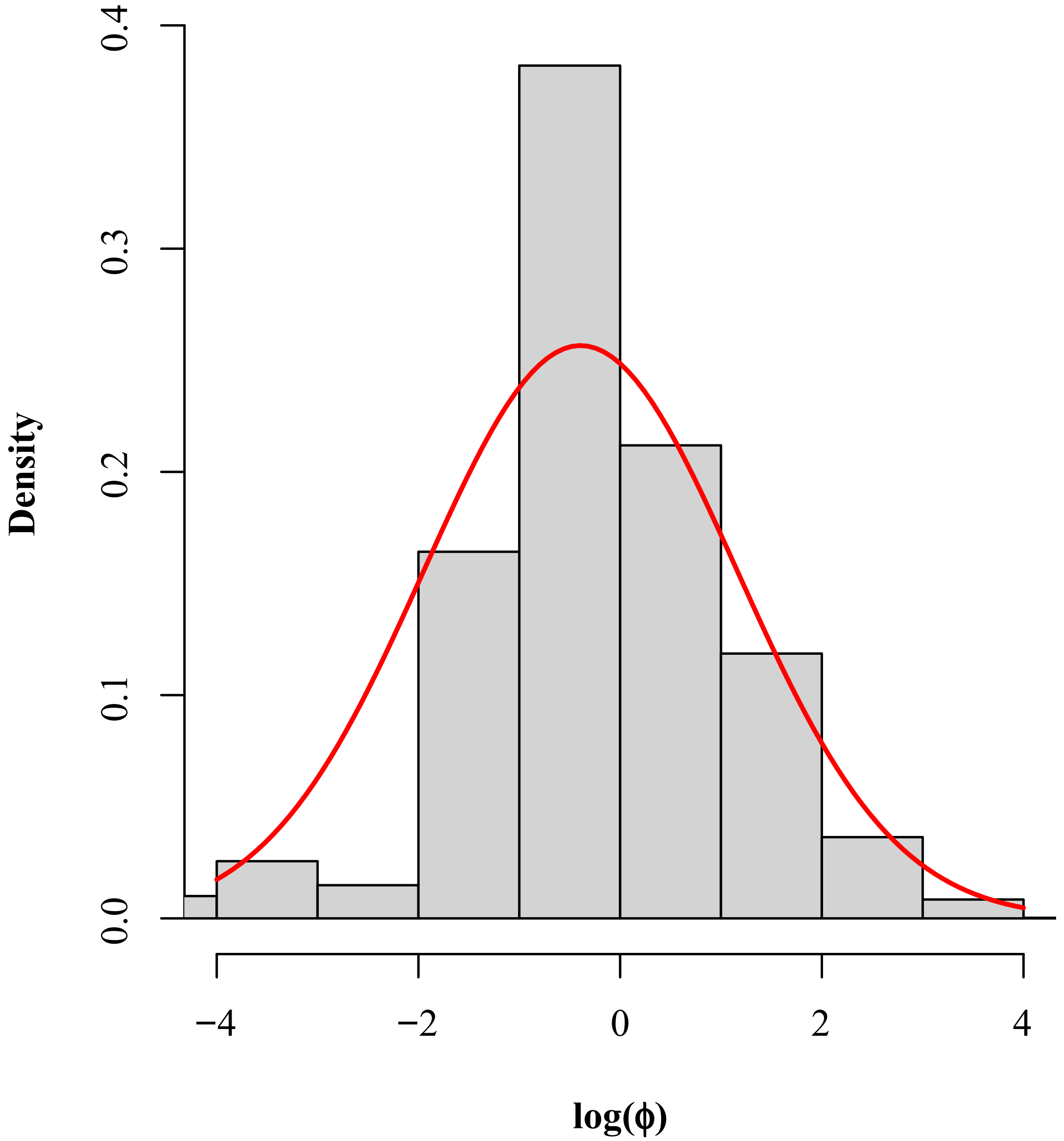


Figure S1. Histogram of log-transformed dispersion estimates in granule nueron from "Human Brain 3K"

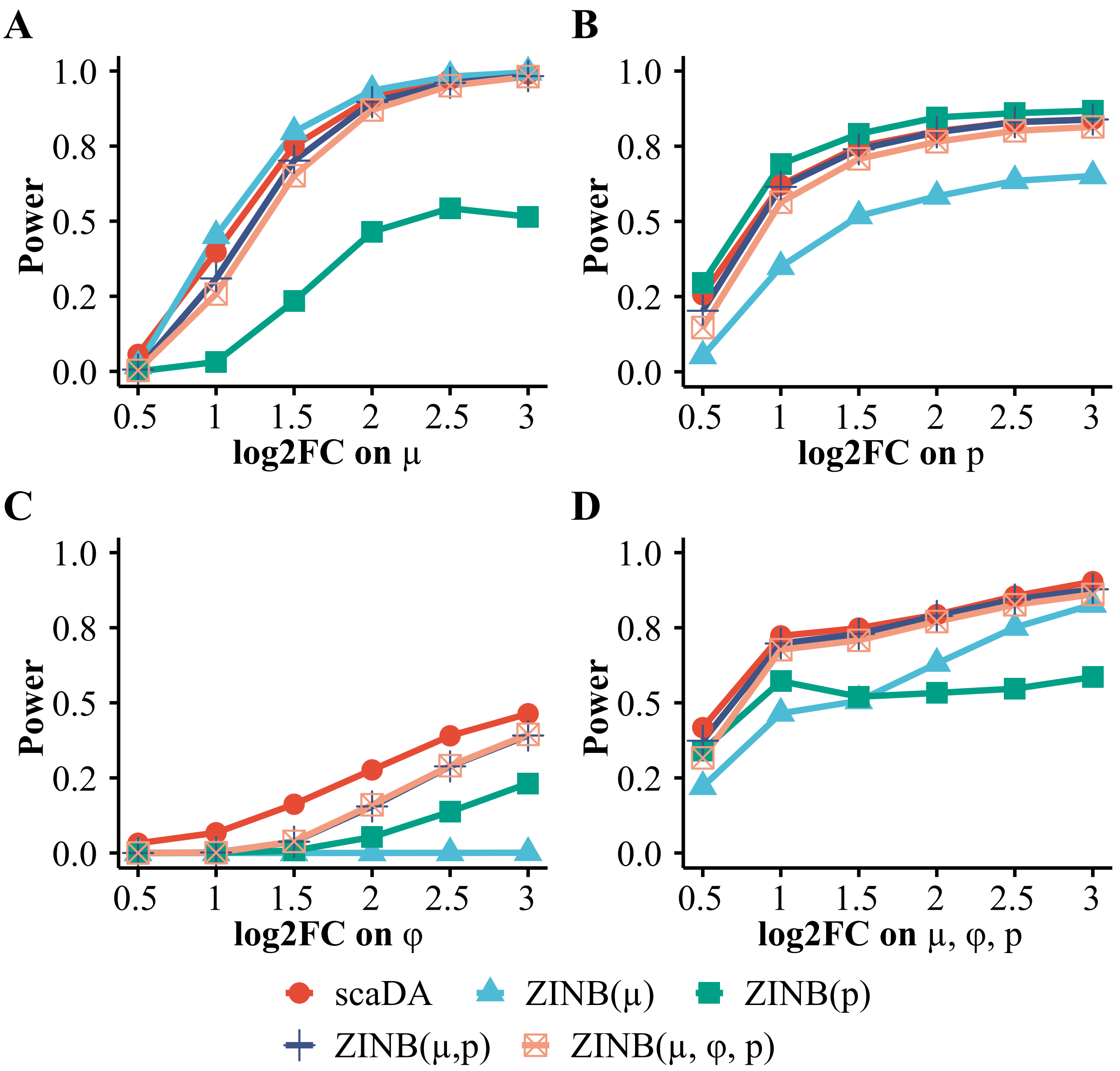


Figure S2. **Power analysis for scaDA and other ZINB-based likelihood ratio tests (Sample size 200 in each group).** Baseline values of three parameters are estimated in the cell type "Granule neuron" from "Human Brain 3K". For DA peaks, we fix the parameter values in Group 1 and vary the parameter values in the Group 2 with different effect size of selected parameters to create power curves. To create parameter values in Group 2, we multiply and divide an effect size in terms of log2 fold change from the baseline values for DA peaks in equal proportion. The log2 fold change of selected parameters is changed from 0.5 to 3.0 with a step size of 0.5. Using the parameter setting in the four scenarios, we assume 20% DA peaks and simulate the read counts based on ZINB for 4000 peaks across 200 cells in each group. **A.** Scenario 1: only mean difference between two groups **B.** Scenario 2: only prevalence between two groups. **C.** Scenario 3: only dispersion difference between two groups. **D.** Scenario 4: difference of all three parameters between two groups.

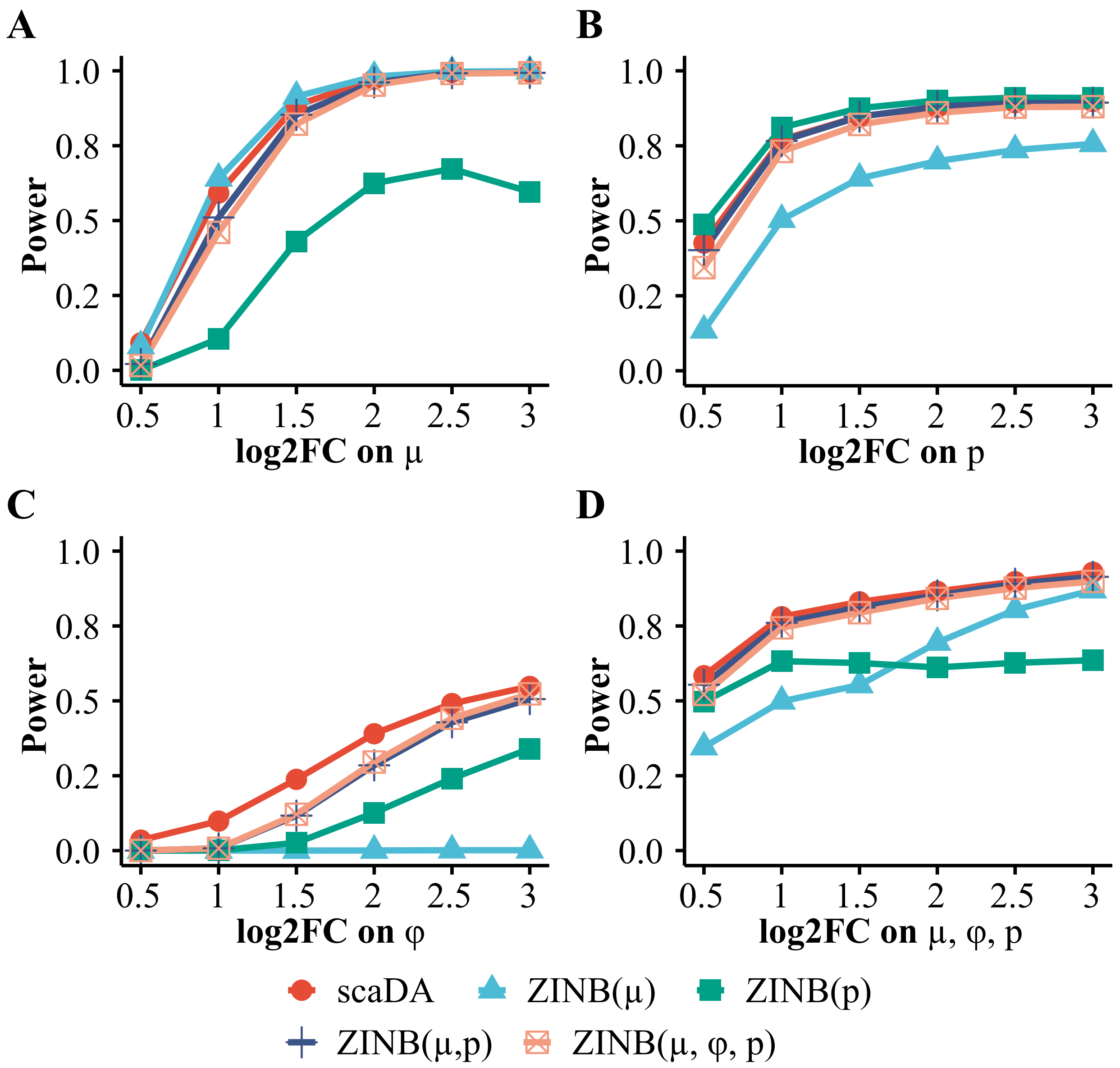


Figure S3. **Power analysis for scaDA and other ZINB-based likelihood ratio tests (Sample size 300 in each group).** Baseline values of three parameters are estimated in the cell type "Granule neuron" from "Human Brain 3K". For DA peaks, we fix the parameter values in Group 1 and vary the parameter values in the Group 2 with different effect size of selected parameters to create power curves. To create parameter values in Group 2, we multiply and divide an effect size in terms of log2 fold change from the baseline values for DA peaks in equal proportion. The log2 fold change of selected parameters is changed from 0.5 to 3.0 with a step size of 0.5. Using the parameter setting in the four scenarios, we assume 20% DA peaks and simulate the read counts based on ZINB for 4000 peaks across 300 cells in each group. **A.** Scenario 1: only mean difference between two groups **B.** Scenario 2: only prevalence between two groups. **C.** Scenario 3: only dispersion difference between two groups. **D.** Scenario 4: difference of all three parameters between two groups.

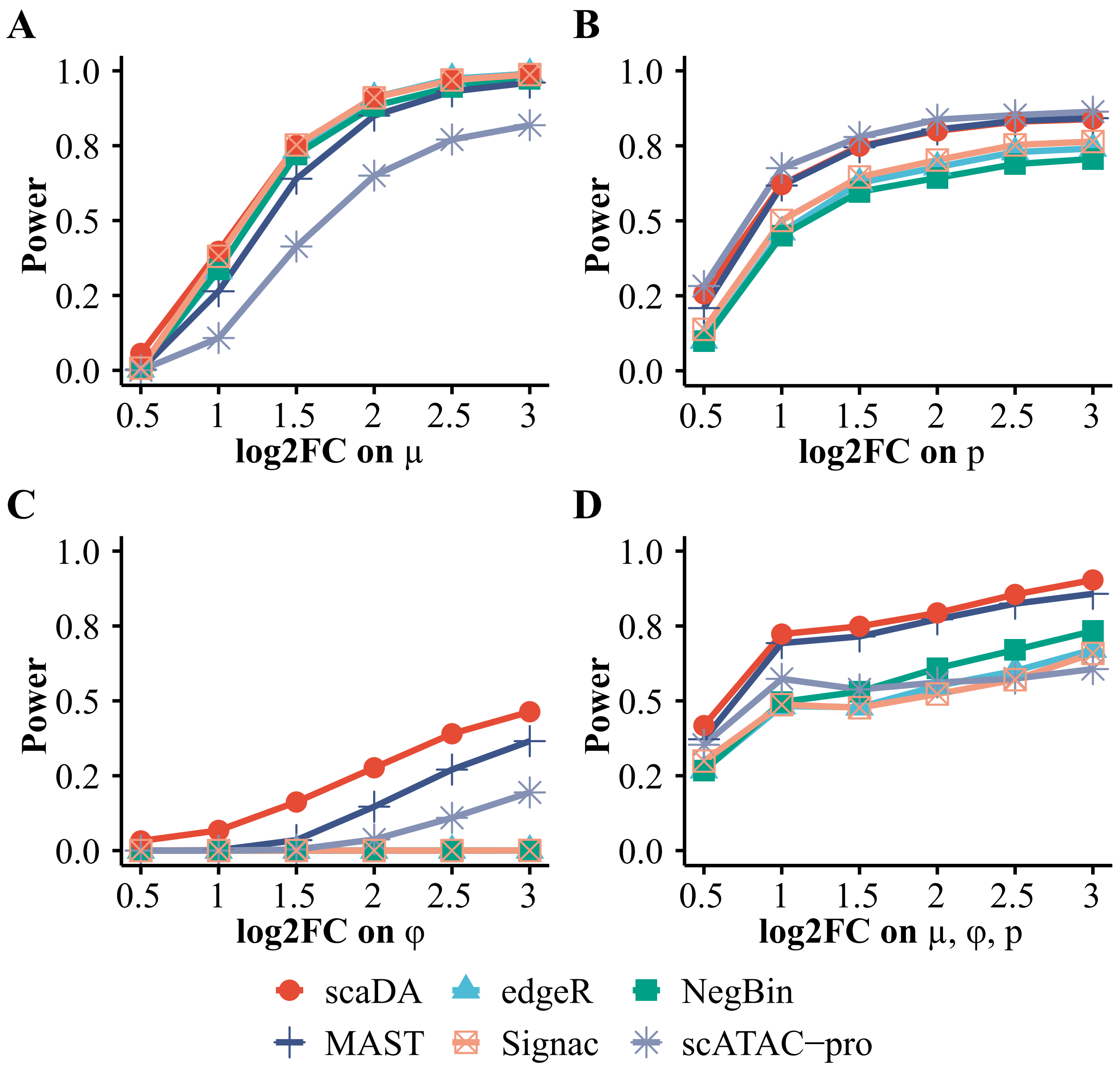


Figure S4. **Power analysis for scaDA and published methods (Sample size 200 in each group).** Baseline values of three parameters are estimated in the cell type "Granule neuron" from "Human Brain 3K". For DA peaks, we fix the parameter values in Group 1 and vary the parameter values in the Group 2 with different effect size of selected parameters to create power curves. To create parameter values in Group 2, we multiply and divide an effect size in terms of log2 fold change from the baseline values for DA peaks in equal proportion. The log2 fold change of selected parameters is changed from 0.5 to 3.0 with a step size of 0.5. Using the parameter setting in the four scenarios, we assume 20% DA peaks and simulate the read counts based on ZINB for 4000 peaks across 200 cells in each group. **A.** Scenario 1: only mean difference between two groups **B.** Scenario 2: only prevalence between two groups. **C.** Scenario 3: only dispersion difference between two groups. **D.** Scenario 4: difference of all three parameters between two groups.

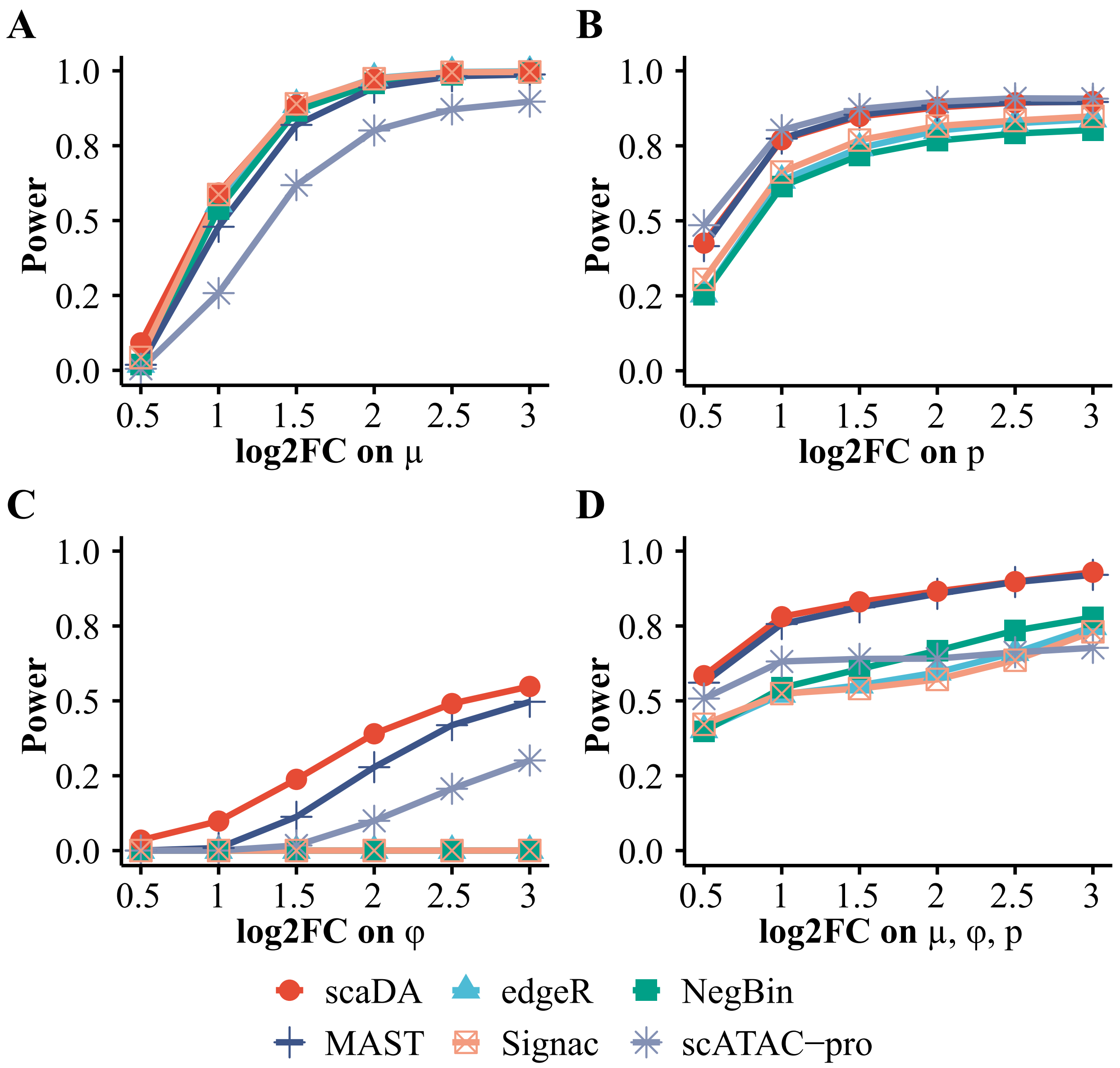


Figure S5. **Power analysis for scaDA and published methods (Sample size 300 in each group).** Baseline values of three parameters are estimated in the cell type "Granule neuron" from "Human Brain 3K". For DA peaks, we fix the parameter values in Group 1 and vary the parameter values in the Group 2 with different effect size of selected parameters to create power curves. To create parameter values in Group 2, we multiply and divide an effect size in terms of log2 fold change from the baseline values for DA peaks in equal proportion. The log2 fold change of selected parameters is changed from 0.5 to 3.0 with a step size of 0.5. Using the parameter setting in the four scenarios, we assume 20% DA peaks and simulate the read counts based on ZINB for 4000 peaks across 300 cells in each group. **A.** Scenario 1: only mean difference between two groups **B.** Scenario 2: only prevalence between two groups. **C.** Scenario 3: only dispersion difference between two groups. **D.** Scenario 4: difference of all three parameters between two groups.

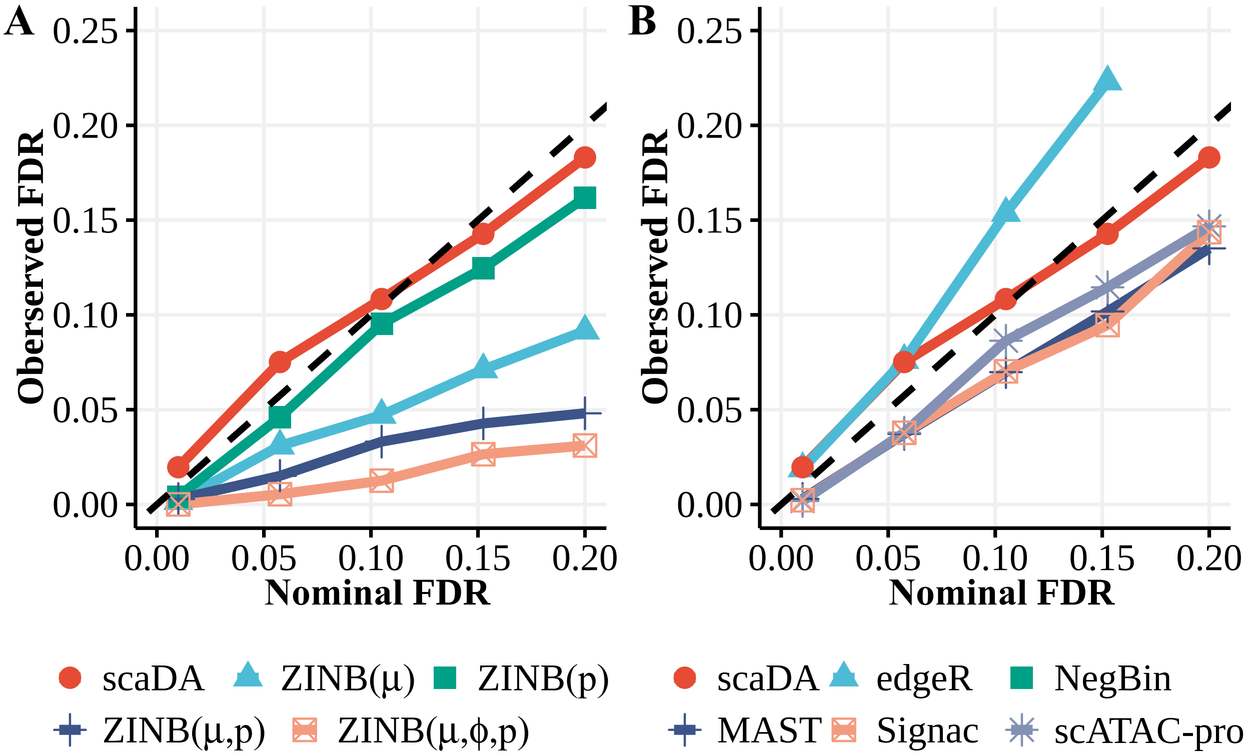


Figure S6. **FDR control analysis (Sample size 300 in each group).**. scaDA is compared to both ZINB-based LRT tests and published methods. Using the same simulation strategy in Scenario 4 (log2FC=3), we assume 20% DA peaks and simulate the read counts based on ZINB for 4000 peaks across 200 cells in each group. The observed FDR is plotted against the nominal FDR level. **A.** scaDA is compared to ZINB-based LRT tests. **B.** scaDA is compared to published methods.

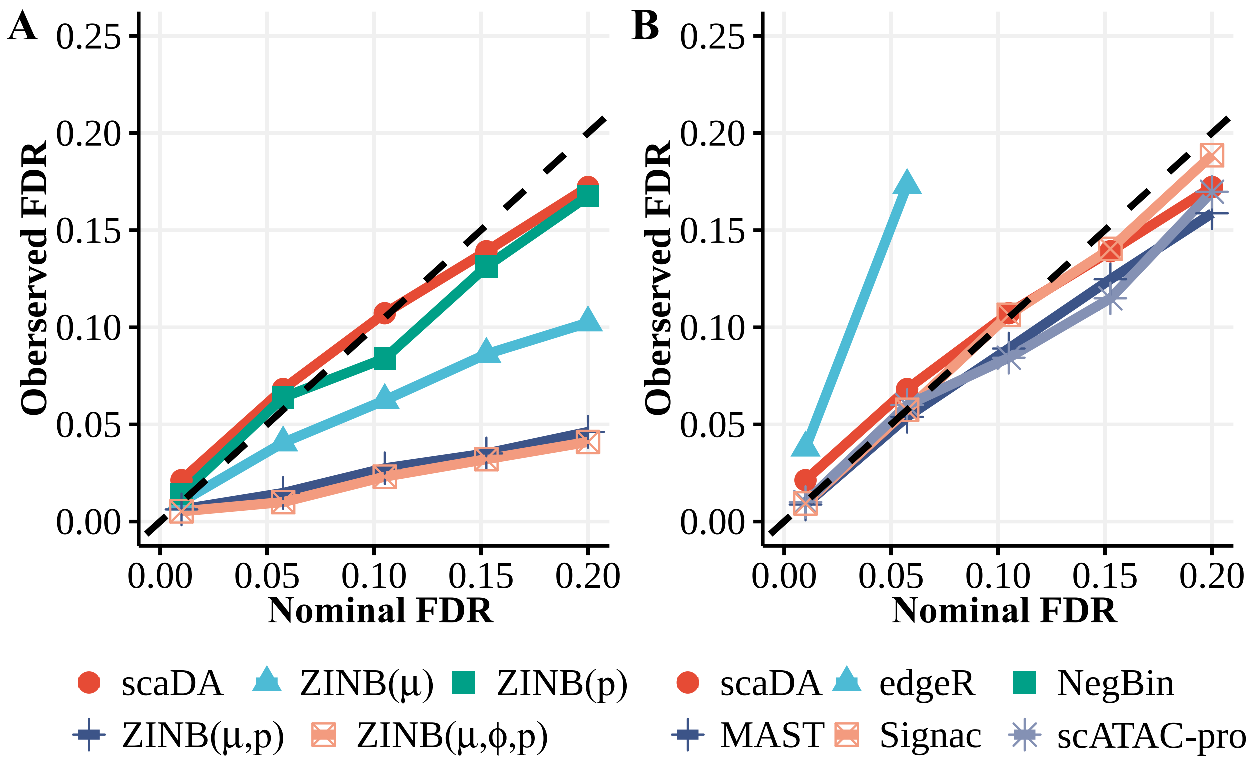


Figure S7. **FDR control analysis (Sample size 300 in each group).**. scaDA is compared to both ZINB-based LRT tests and published methods. Using the same simulation strategy in Scenario 4 (log2FC=3), we assume 20% DA peaks and simulate the read counts based on ZINB for 4000 peaks across 300 cells in each group. The observed FDR is plotted against the nominal FDR level. **A.** scaDA is compared to ZINB-based LRT tests. **B.** scaDA is compared to published methods.

#### Real data analysis

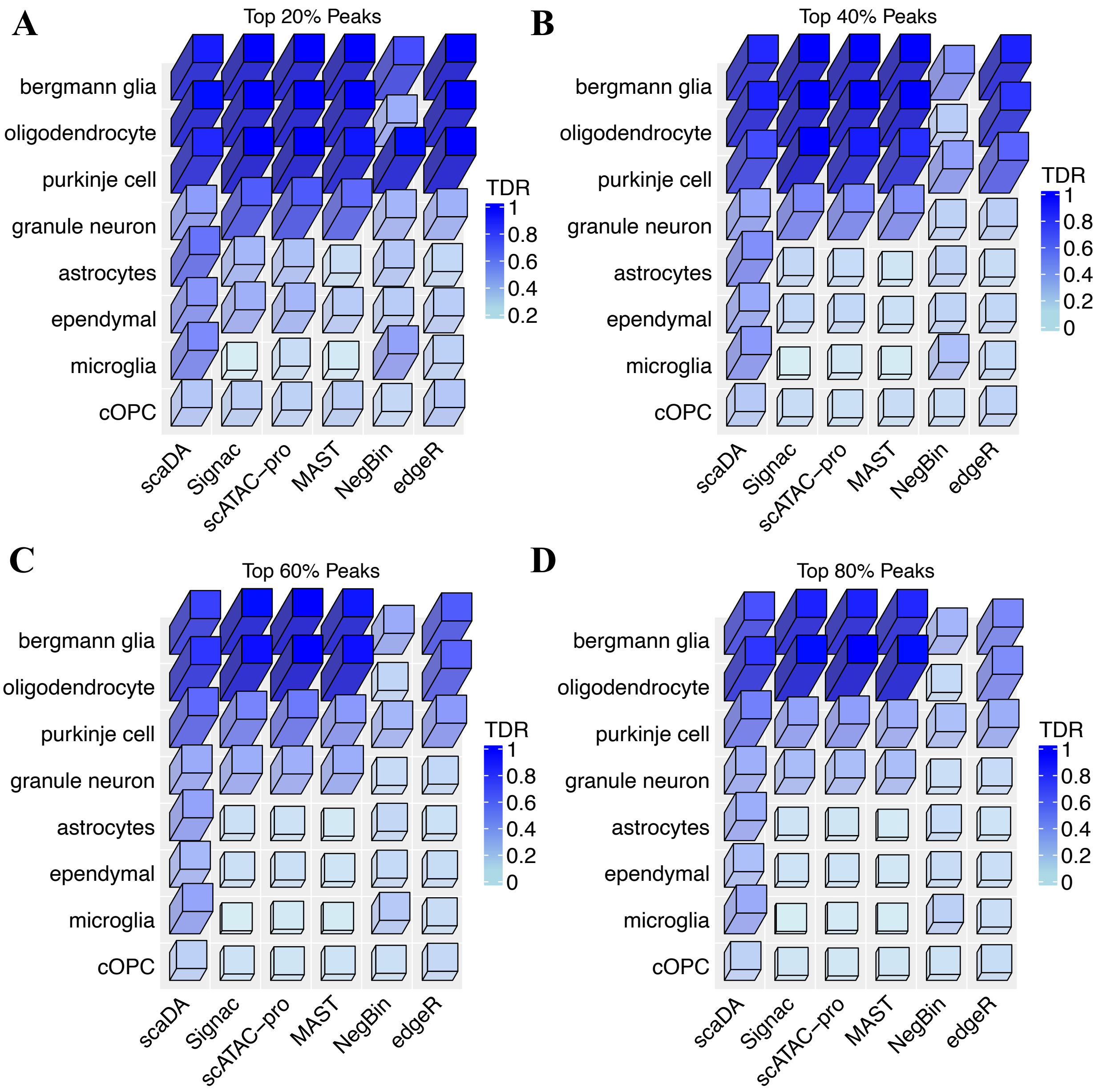


Figure S8. Human Brain 3K: TDR across 8 cell types for all methods at different levels of top percentages (20%, 40%, 60%, 80%)

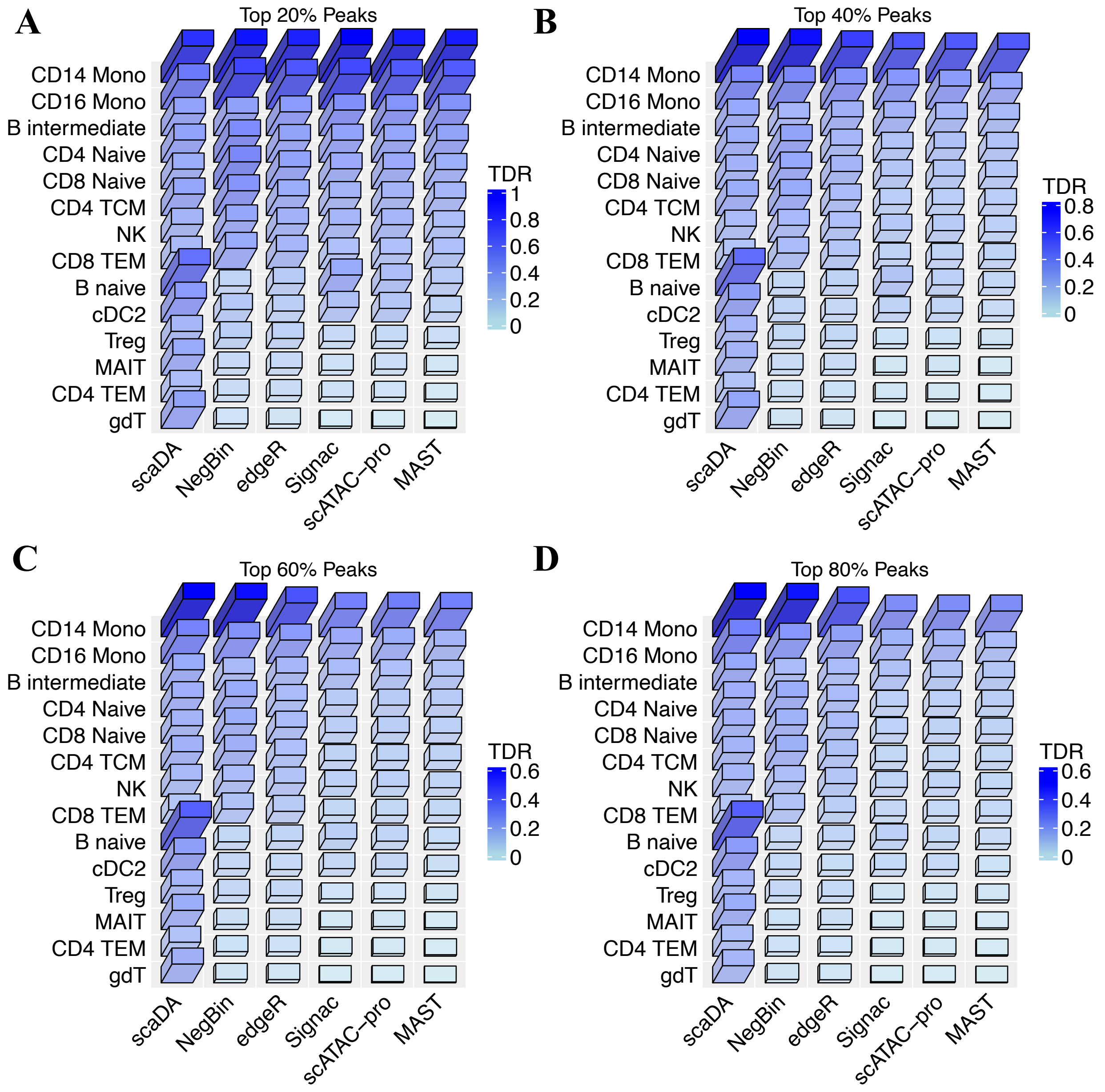


Figure S9. Human PBMC 10K: TDR across 14 cell types for all methods at different levels of top percentages (20%, 40%, 60%, 80%)

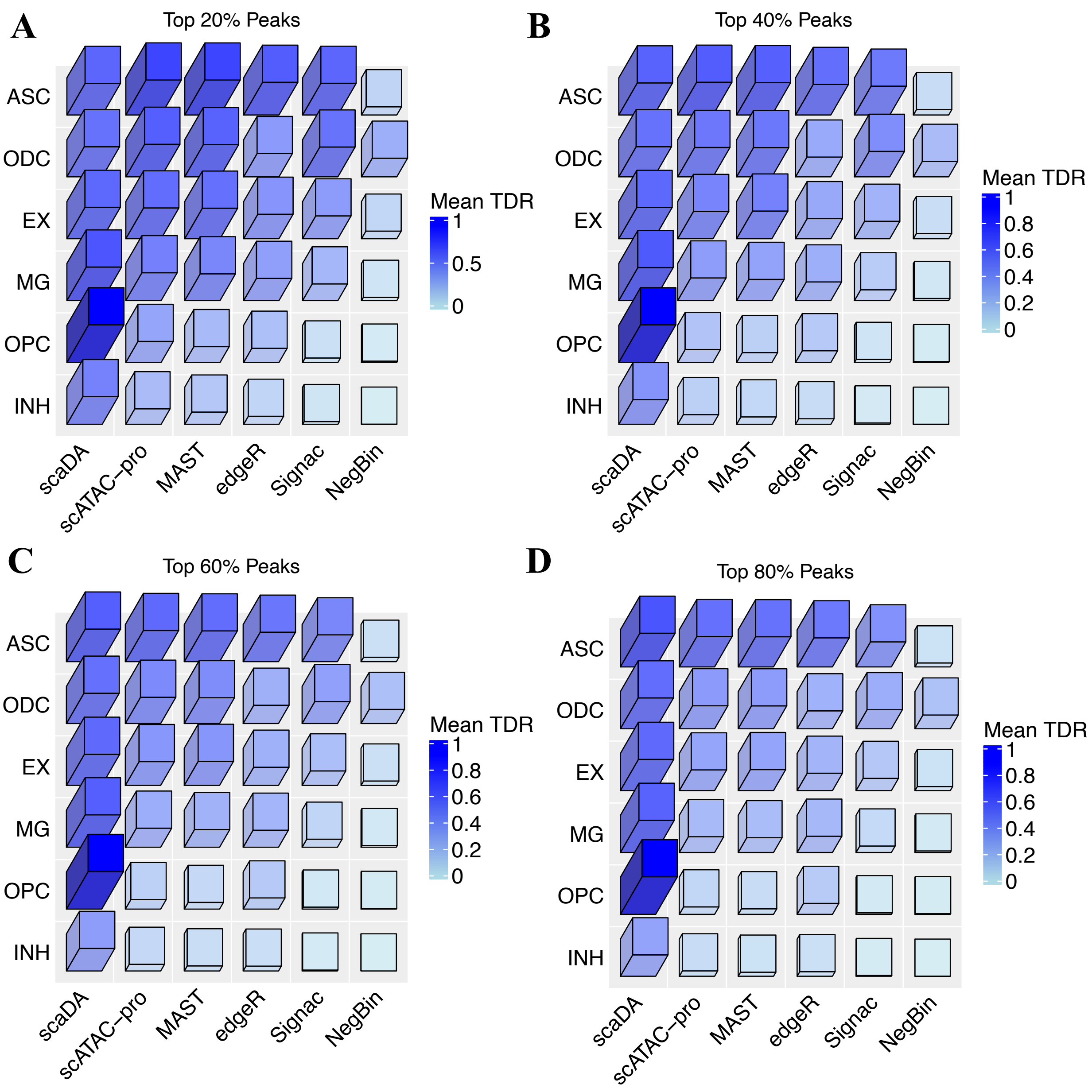


Figure S10. Human AD: TDR across 6 cell types for all methods at different levels of top percentages (20%, 40%, 60%, 80%)

### Supplemental Tables

#### Human Brain 3K

| celltype | cellnum | cell Proportion |
|---|---|---|
| granule neuron | 636 | 22% |
| oligodendrocyte | 552 | 20% |
| cOPC | 365 | 13% |
| bergmann glia | 344 | 12% |
| ependymal | 204 | 7% |
| purkinje cell | 144 | 5% |
| astrocytes | 125 | 4% |
| microglia | 104 | 4% |
| GCP | 93 | 3% |
| brainstem | 73 | 3% |
| MLI | 69 | 2% |
| rhombic lip | 63 | 2% |
| iCN | 29 | 1% |
| UBC | 29 | 1% |

Table S1. Human Brain 3K: Cell types, cell numbers and cell type proportion

| Top Peaks | scaDA | Signac | scATAC-pro | MAST | NegBin | edgeR |
|---|---|---|---|---|---|---|
| 20% | 0.81 | 0.72 | 0.72 | 0.67 | 0.64 | 0.67 |
| 40% | 0.73 | 0.58 | 0.58 | 0.56 | 0.42 | 0.54 |
| 60% | 0.66 | 0.48 | 0.49 | 0.46 | 0.32 | 0.44 |
| 80% | 0.59 | 0.42 | 0.42 | 0.40 | 0.26 | 0.35 |
| 100% | 0.51 | 0.34 | 0.34 | 0.33 | 0.22 | 0.28 |

Table S2. Human Brain 3K: Mean of TDR across all cell types at different levels of top percentages

| Top Peaks | scaDA | Signac | scATAC-pro | MAST | NegBin | edgeR |
|---|---|---|---|---|---|---|
| 20% | 0.03 | 0.09 | 0.08 | 0.10 | 0.05 | 0.08 |
| 40% | 0.05 | 0.15 | 0.15 | 0.16 | 0.03 | 0.11 |
| 60% | 0.05 | 0.15 | 0.15 | 0.15 | 0.02 | 0.09 |
| 80% | 0.05 | 0.14 | 0.14 | 0.14 | 0.01 | 0.06 |
| 100% | 0.03 | 0.09 | 0.09 | 0.10 | 0.01 | 0.03 |

Table S3. Human Brain 3K: Variance of TDR across all cell types at different levels of top percentages

| | scaDA | Signac | scATAC-pro | MAST | NegBin | edgeR |
|---|---|---|---|---|---|---|
| Rank 1 | 6 | 2 | 0 | 1 | 0 | 0 |
| Rank 2 | 0 | 1 | 2 | 0 | 3 | 1 |
| Rank 3 | 0 | 1 | 1 | 2 | 1 | 3 |
| Rank 4 | 2 | 3 | 2 | 0 | 0 | 1 |
| Rank 5 | 0 | 0 | 2 | 3 | 0 | 3 |
| Rank 6 | 0 | 1 | 1 | 2 | 4 | 0 |

Table S4. Human Brain 3K: Rank of all methods by power

#### Human PBMC 10K

| celltype | cellnum | cellprop |
|---|---|---|
| CD4 TCM | 2303 | 27% |
| CD14 Mono | 2124 | 25% |
| CD4 Naive | 858 | 10% |
| CD8 TEM | 801 | 9% |
| CD8 Naive | 515 | 6% |
| NK | 336 | 4% |
| B intermediate | 318 | 4% |
| CD16 Mono | 266 | 3% |
| Treg | 187 | 2% |
| CD4 TEM | 154 | 2% |
| MAIT | 144 | 2% |
| cDC2 | 119 | 1% |
| B naive | 112 | 1% |
| gdT | 111 | 1% |
| CD8 TCM | 97 | 1% |
| B memory | 85 | 1% |
| pDC | 43 | 0% |
| CD4 CTL | 24 | 0% |
| NK_CD56bright | 21 | 0% |
| Plasmablast | 16 | 0% |

Table S5. Human PBMC 10K: Cell types, cell numbers and cell type proportion

| Top Peaks | scaDA | NegBin | edgeR | Signac | scATAC-pro | MAST |
|---|---|---|---|---|---|---|
| 20% | 0.57 | 0.49 | 0.45 | 0.45 | 0.43 | 0.40 |
| 40% | 0.43 | 0.31 | 0.29 | 0.25 | 0.24 | 0.22 |
| 60% | 0.34 | 0.24 | 0.21 | 0.17 | 0.16 | 0.15 |
| 80% | 0.29 | 0.19 | 0.17 | 0.12 | 0.12 | 0.11 |
| 100% | 0.24 | 0.16 | 0.14 | 0.10 | 0.10 | 0.09 |

Table S6. Human PBMC 10K: Mean of TDR across all cell types at different levels of top percentages

| Top Peaks | scaDA | NegBin | edgeR | Signac | scATAC-pro | MAST |
|---|---|---|---|---|---|---|
| 20% | 0.02 | 0.07 | 0.06 | 0.07 | 0.07 | 0.08 |
| 40% | 0.02 | 0.04 | 0.03 | 0.03 | 0.03 | 0.03 |
| 60% | 0.01 | 0.02 | 0.02 | 0.01 | 0.01 | 0.01 |
| 80% | 0.01 | 0.02 | 0.01 | 0.01 | 0.01 | 0.01 |
| 100% | 0.01 | 0.01 | 0.01 | 0.01 | 0.01 | 0.01 |

Table S7. Human PBMC 3K: Variance of TDR across all cell types at different levels of top percentages

| | scaDA | NegBin | edgeR | Signac | scATAC-pro | MAST |
|---|---|---|---|---|---|---|
| Rank 1 | 9 | 5 | 0 | 0 | 0 | 0 |
| Rank 2 | 5 | 8 | 1 | 1 | 0 | 0 |
| Rank 3 | 0 | 0 | 12 | 1 | 0 | 0 |
| Rank 4 | 0 | 1 | 0 | 10 | 6 | 4 |
| Rank 5 | 0 | 0 | 1 | 2 | 6 | 0 |
| Rank 6 | 0 | 0 | 0 | 0 | 2 | 10 |

Table S8. Human Brain 3K: Rank of all methods by power

#### Human AD

| Batch | Control | Case |
|---|---|---|
| 1 | Sample-96 | Sample-43 |
| 1 | Sample-100 | Sample-45 |
| 2 | Sample-82 | Sample-46 |
| 2 | Sample-66 | Sample-40 |
| 2 | Sample-101 | Sample-27 |
| 2 | | Sample-50 |
| 2 | | Sample-22 |
| 3 | Sample-90 | Sample-37 |
| 3 | Sample-52 | Sample-33 |
| 3 | Sample-58 | Sample-47 |
| 3 | | Sample-17 |
| 3 | | Sample-19 |

Table S9. Human AD: experimental design

| Top peaks | scaDA | scATAC-pro | MAST | edgeR | Signac | NegBin |
|---|---|---|---|---|---|---|
| 20% | 0.77 | 0.65 | 0.62 | 0.51 | 0.45 | 0.16 |
| 40% | 0.75 | 0.54 | 0.52 | 0.43 | 0.35 | 0.12 |
| 60% | 0.73 | 0.46 | 0.44 | 0.39 | 0.30 | 0.10 |
| 80% | 0.71 | 0.40 | 0.38 | 0.37 | 0.25 | 0.09 |
| 100% | 0.69 | 0.34 | 0.33 | 0.35 | 0.21 | 0.09 |

Table S10. Human AD: Mean of TDR across all cell types and pairwise comparison at different levels of top percentages

| Top peaks | scaDA | scATAC-pro | MAST | edgeR | Signac | NegBin |
|---|---|---|---|---|---|---|
| 20% | 0.10 | 0.17 | 0.17 | 0.16 | 0.19 | 0.07 |
| 40% | 0.10 | 0.17 | 0.17 | 0.14 | 0.17 | 0.05 |
| 60% | 0.10 | 0.16 | 0.16 | 0.13 | 0.14 | 0.04 |
| 80% | 0.10 | 0.14 | 0.14 | 0.12 | 0.12 | 0.04 |
| 100% | 0.09 | 0.11 | 0.11 | 0.11 | 0.09 | 0.04 |

Table S11. Human AD: Variance of TDR across all cell types and pairwise comparison at different levels of top percentages

| | scaDA | scATAC-pro | MAST | edgeR | Signac | NegBin |
|---|---|---|---|---|---|---|
| Rank 1 | 6 | 0 | 0 | 0 | 0 | 0 |
| Rank 2 | 0 | 3 | 1 | 2 | 0 | 0 |
| Rank 3 | 0 | 3 | 2 | 1 | 0 | 0 |
| Rank 4 | 0 | 0 | 3 | 3 | 0 | 0 |
| Rank 5 | 0 | 0 | 0 | 0 | 6 | 0 |
| Rank 6 | 0 | 0 | 0 | 0 | 0 | 6 |

Table S12. Human AD: Rank of all methods by mean power
